# ASPEN: Robust detection of allelic dynamics in single cell RNA-seq

**DOI:** 10.1101/2025.04.16.649227

**Authors:** Veronika Petrova, Muqing Niu, Thomas Vierbuchen, Emily S Wong

## Abstract

Single-cell RNA-seq data from F1 hybrids provides a unique framework for dissecting complex regulatory phenomena, but allelic measurements are limited by technical noise. Here, we present ASPEN, a statistical method for modeling allelic mean and variance in single-cell transcriptomic data from F1 hybrids. ASPEN uses a sensitive mapping pipeline and adaptive shrinkage to distinguish allelic imbalance and variance in single cells. Through extensive simulation based on sparse droplet-based single-cell data, ASPEN demonstrates improved sensitivity and control of false discoveries compared to existing approaches. Applied to mouse brain organoids and T cells, ASPEN identifies genes with incomplete X inactivation, stochastic monoallelic expression, and significant deviations in allelic variance. This reveals reduced variance in essential cellular pathways, and increased variance in neurodevelopmental and immune-specific genes.

## Background

F1 hybrids, the offsprings of inbred parental strains, provide a powerful model for identifying the genetic influences that impact gene regulation. By combining a controlled genetic background within a shared nuclear environment, F1 hybrids model the contributions of cis- and trans-regulatory variation to changes in gene regulation. These hybrids have been used to uncover core principles of transcriptional regulation and epigenetic processes, including parent-of-origin effects ^1–3^, X-chromosome inactivation^4,5^, cis regulatory evolution ^6,7^, and the molecular basis of hybrid vigor ^9,2^. The approach, with high-throughput sequencing, has been applied across taxa, from model mammals (e.g., mice ^8,9^) and insects (e.g., *Drosophila* ^10–12^) to plants (e.g., *Arabidopsis* ^13–15^) and fungi ^16–18^.

Study of cis-regulatory mechanisms in F1 hybrids combined with single cell genomics allows resolution of phenomena such as transcriptional bursting ^19,20^, incomplete X inactivation ^21,22^, and cis-regulation across lineage differentiation ^2^□. Several analytical frameworks have been developed to assess allelic changes using single-cell RNA-seq data. These include Airpart, which uses a hierarchical Bayesian model to estimate cell type-specific allelic imbalances^23^; scDALI, which implements a generalized linear mixed model to detect variations in ASE across continuous trajectories^24^; and SCALE, which compares transcriptional kinetics between alleles within a hierarchical Bayesian framework^19^. Methods for allelic quantification may also focus on natural populations. For example, DAESC uses random effects modelling to study allelic usage in humans ^25^. These tools have advanced the field, yet challenges remain, particularly due to data sparsity in single cell data, and the detection subtle regulatory shifts and changes across cell states.

To address these limitations, we introduce ASPEN, a flexible statistical method for modeling allelic imbalance and variance in single-cell transcriptomic data. ASPEN uses a sensitive allele-specific quantification pipeline with a moderated beta-binomial model that incorporates adaptive shrinkage to stabilize dispersion estimates allowing the detection of allele-specific expression patterns using scRNA-seq. Validated against simulated and empirical datasets, ASPEN demonstrates a 30% increase in sensitivity over existing approaches for single cell allelic imbalance detection. Applying ASPEN to mouse brain organoids and T cells, we uncover dynamic cis-regulatory changes during cellular differentiation, stochastic monoallelic expression, and incomplete dosage compensation in X-linked genes.

## Results and Discussion

### Regulatory control, not technical noise, underlies low allelic variation

Analyses of single-cell allele-specific expression (scASE) data, particularly the high sparsity of droplet-based data such as 10x Chromium, is challenging due to high technical noise which can obscure true biological signal ^26^. To address this, we implemented a sensitive mapping pipeline with a moderated beta-binomial model that features conditional shrinkage to accurately detect allele-specific gene regulation using single-cell data (**Fig. 1**).

**Figure 1.**
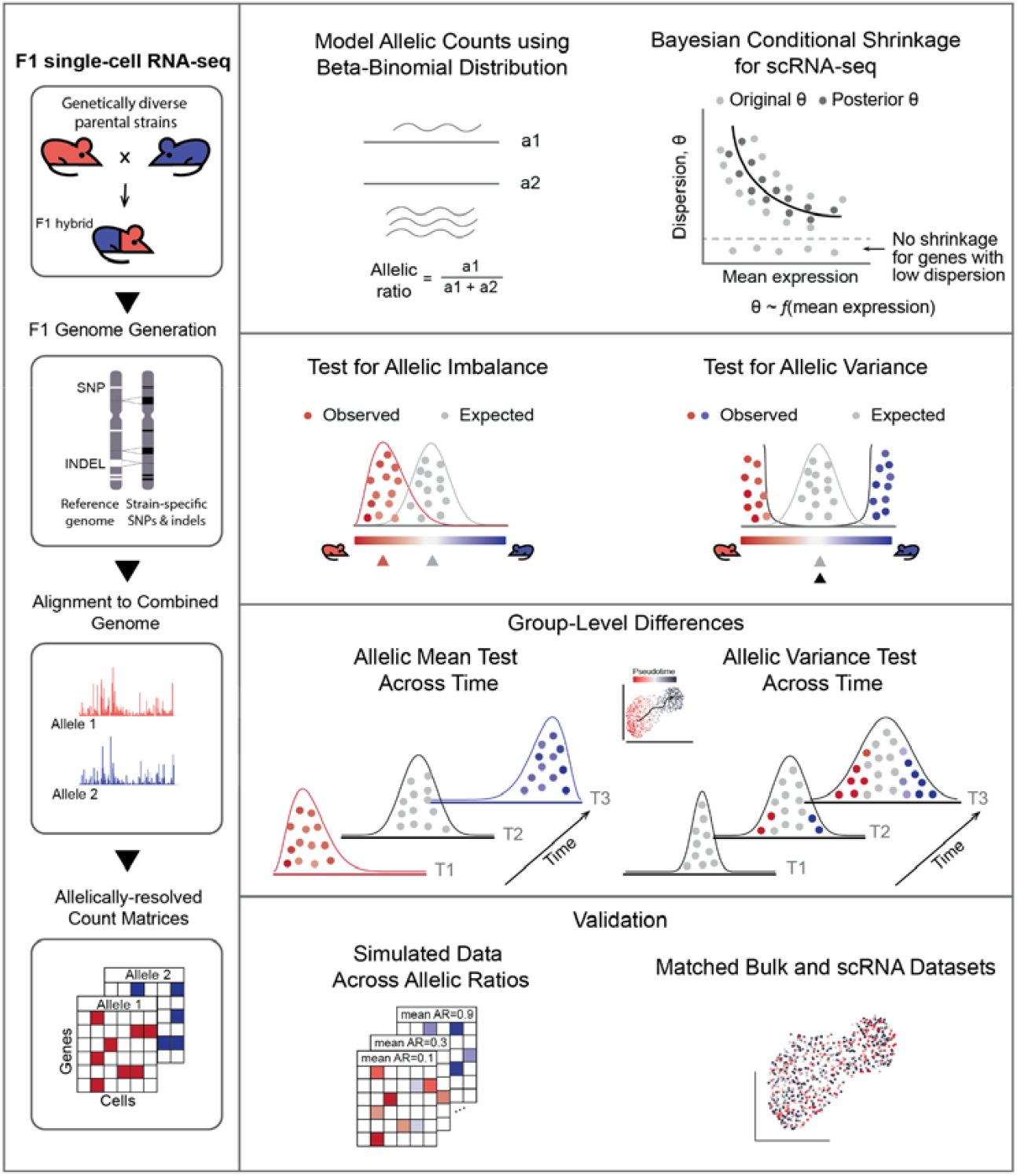
Overview of ASPEN. LHS: Raw sequencing reads are mapped to a combined genome created by concatenating the two parental genomes. The C57BL/6J (GRCm38) genome serves as the reference, while strain-specific genomes are produced by integrating variant information (SNPs and indels) into the C57BL/6J genome. RHS: ASPEN employs the Bayesian shrinkage method to stabilize dispersion estimates across genes with varying expression levels. Using weighted log-likelihood, original dispersion estimates are shrunk toward the common trend dispersion — the expected level of variation for genes exhibiting similar expression levels. The posterior (shrunken) dispersion estimates facilitate the detection of state-specific and differential changes in the allelic ratio distribution.

We first implemented a stringent read-mapping pipeline that integrates both single nucleotide polymorphisms (SNPs) and small insertions/deletions (indels) to align reads against custom F1 hybrid mouse genome assemblies. We created custom assemblies for common F1 crosses between the reference *C57BL/6J* strain and *CAST/Ei, MOLF/Ei, PWK/Ph, and SPRET/Ei* from the Hybrid Mouse Diversity Panel. By incorporating both SNPs and indels in the genome references, read mapping was improved by ∼10% compared to using SNPs alone (**Supp Fig. 1**). We only retained uniquely mapped reads for unambiguous allelic assignment as the incorporation of multi-mapping reads to increase read coverage reduced the number of allelic imbalance genes identified (**Supplementary Results**). Across four widely used mouse hybrid crosses, this strategy enabled the resolution of 20–38% of total sequencing reads. Due to genome quality, most aligned reads mapped to the reference genome (52-56%). ASPEN’s allelic imbalance test accounts for this mapping bias by adjusting the expected background ratio.

To improve statistical power of resolving allelic counts while accounting for data sparsity, we used a moderated beta-binomial modeling framework with empirical Bayes shrinkage of dispersion parameters. Allelic variance was parameterized as dispersion by quantifying the deviation around the beta-binomial mean allele ratio. Shrinkage was applied across the transcriptome, stabilizing dispersion estimates by borrowing information across genes while still accommodating gene-specific departures from the global trend reducing technical noise ^27,28^. This approach is conceptually similar to that used in the differential gene expression analyses of bulk RNA-seq data 29,30.

We observed a set of low variance outlier genes that clustered distinctly below the modelled relationship between allelic dispersion and gene expression, despite meeting minimal read coverage threshold (**Fig. 2A**). We hypothesized that these genes, which exhibit consistently low allelic dispersion, represent loci under stringent cis-regulatory control to maintain stable expression across cells. In such cases, overshrinkage could artificially inflate dispersion estimates and mask biologically relevant signals. To assess this, we compared allelic dispersion profiles between single-cell and bulk RNA-seq datasets generated from activated CD8+ T cells (day 7 post-infection) in *C57BL/6J* × *SPRET/Ei* F1 hybrids ^31,32^. Genes with low dispersion in single-cell data (θ < 0.005) maintained highly stable allelic ratios and exhibited robust expression in bulk RNA-seq (Wilcoxon Z = –8.23, *p* < 2.2 × 10□^1^ □; **Fig. 2A**).

**Figure 2.**
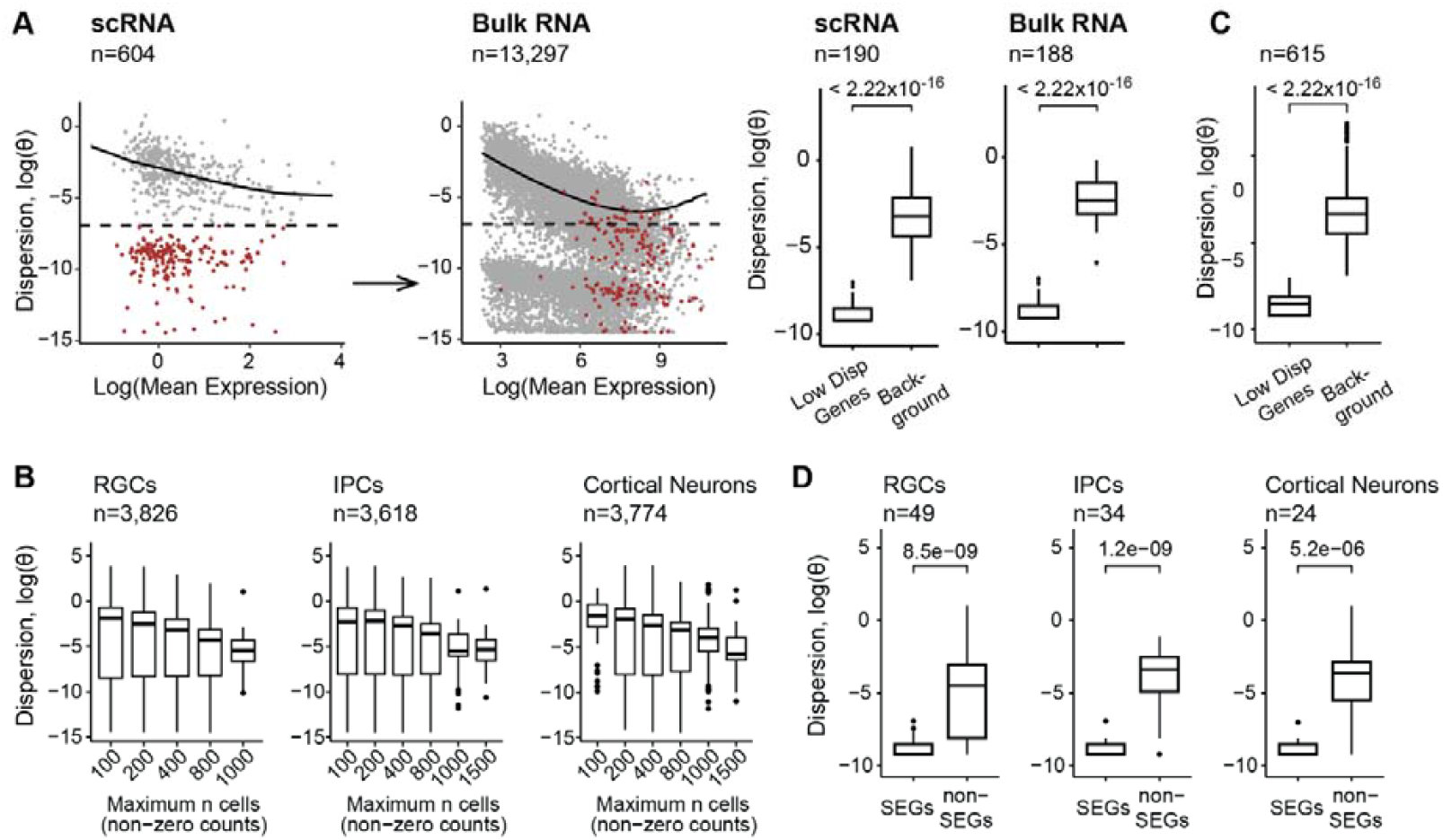
ASPEN performs adaptive dispersion shrinkage tailored for single-cell ASE analysis. **(A)** Evaluating the concordance of low dispersion for genes in matched bulk and single-cell data. Genes with θ < 0.001 in the scRNA counts are marked in red (left panel). The same genes are indicated in red based on the dispersion estimation in the bulk RNA-seq data (right panel). Comparisons were conducted using T-cell data collected on day 7 after LCMV infection. Boxplots illustrate the mean dispersion estimation between low-dispersed genes and the background, a set of genes matched by gene expression (two-sided Wilcoxon rank-sum test). **(B)** Boxplots showing a reduction in variation levels as the number of available (non-zero count) cells increases: RGCs – H-statistic = 216.55, p < 2.22 × 10^-16^, IPCs – H-statistic = 177.42, p < 2.22 × 10^-16^, Cortical neurons –H-statistic = 160.54, p < 2.22 × 10^-16^, Kruskal-Wallis rank sum test). **(C)** Low variation levels are not driven by gene expression or limited number of cells. Dispersion estimation comparison between genes with low dispersion (θ < 0.001) and background, genes matched by cells and reads (two-sided Wilcoxon rank-sum test). **(D)** SEGs demonstrate persistent low dispersion levels when compared with non-SEGs (two-sided Wilcoxon rank-sum test). Background (non-SEGs) is matched to SEGs by gene expression and number of detected cells.

Functional enrichment analysis revealed the under dispersed genes were significantly enriched in housekeeping processes (FDR < 0.1; **Supp. Fig. 2A**), suggesting these pathways may be subject to stronger constraint and tightly coordinated expression. To assess whether the observed low dispersion was the result of sampling noise from low cell numbers and low expression, we randomly subsampled cells to match the distribution of average read counts and the number of cells with non-zero expression in low-dispersion genes (**Fig. 2B**). In contrast, the matched background genes exhibited much greater dispersion, indicating that low dispersion observed was driven by biological effects rather than sampling noise (*p* < 2.22 × 10□^16^, two-sided Wilcoxon test, **Fig. 2C**).

We further compared our dispersion estimates to a list of stably expressed genes (SEGs) identified through integrative analyses across multiple tissues and conditions ^33^, providing an independent benchmark for assessing gene expression stability. Genes classified as SEGs had lower allelic dispersion in our single-cell dataset compared to non-SEGs at matched expression level and cell coverage (median Δθ = 0.2, Wilcoxon p ≤ 5.2 × 10□^6^; **Fig. 2D, Supp Fig. 2B**). SEGs were also generally more enriched among our low-variance genes (Fisher’s exact, OR = 1.3, *p =* 0.04), suggesting these genes were under stronger cis-regulatory control.

In summary, we identified a set of genes with low allelic variances in scRNA-seq datasets, enriched for core housekeeping functions. Their variance remained low even at high expression levels, consistent with stable regulatory control.

### ASPEN robustly detects allelic imbalance in single-cell data

Our above analyses suggest genes with low dispersion likely reflects biological constraint. Hence, ASPEN does not apply shrinkage to those genes with low dispersion. We reasoned that for those genes, shrinkage would inflate their dispersion and reduce the statistical power to detect imbalance (**Fig. 3A, Methods**). Following dispersion parameter estimation, to test for allelic imbalance ASPEN uses a likelihood ratio test to assess the fit of the data against two models. Our null model assumes a fixed mean corresponding to empirically determined allelic balance, while the alternative model estimates the mean allelic ratio directly from the data using maximum likelihood estimation (MLE).

**Figure 3.**
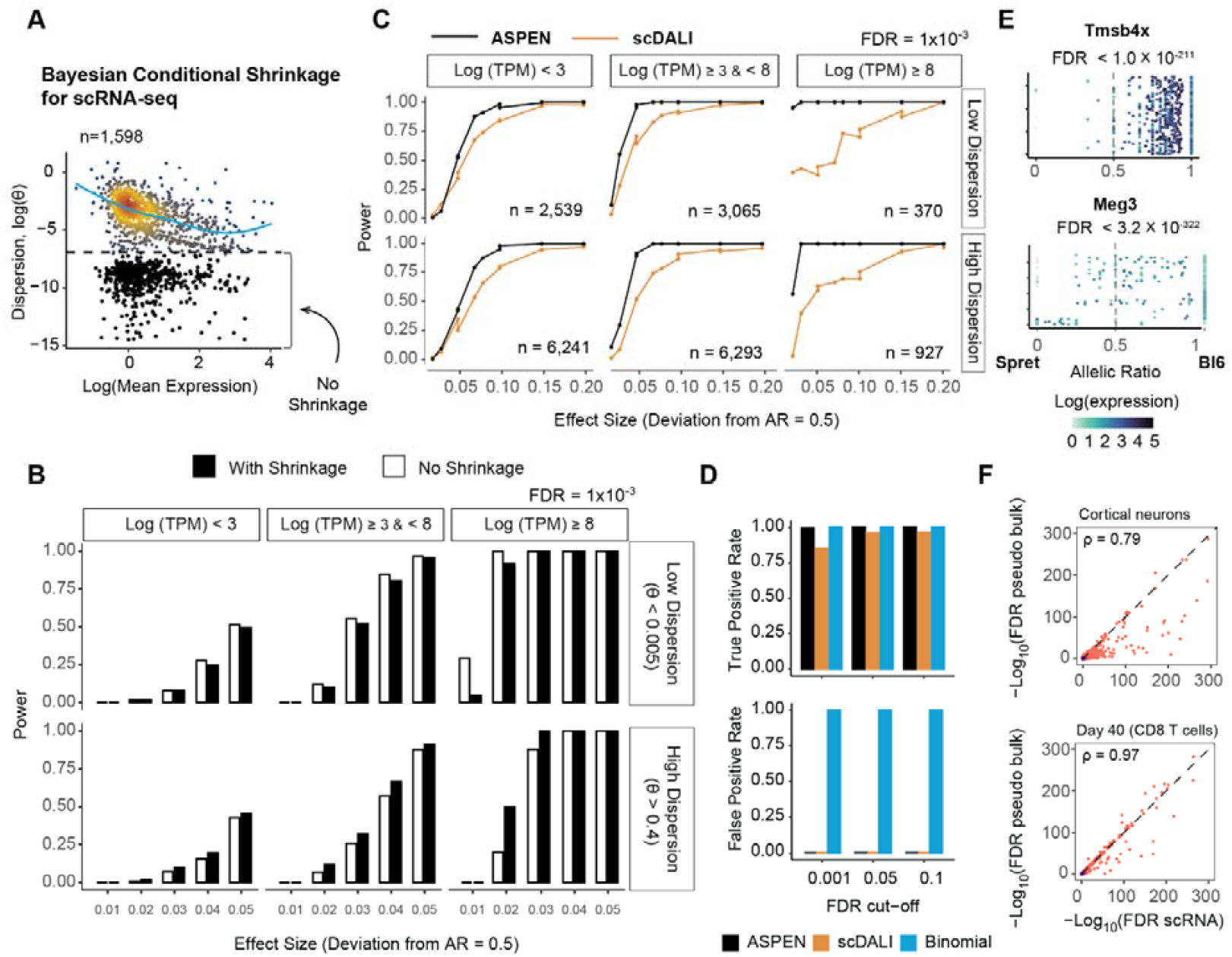
Evaluation and comparison of ASPEN’s performance in detecting ASE imbalance. **(A)** Modeling the relationship between allelic ratio dispersion and mean gene expression revealed a population of genes with persistent low allelic dispersion in T-cell data from male Bl6 × Spret, combining all cell states. The dashed line separates genes with low variation (in black), which are excluded from the shrinkage procedure. **(B)** Evaluating Bayesian (shrinkage applied to all genes) and non-Bayesian (no shrinkage applied) strategies for testing allelic imbalance. Using simulated data, power was estimated as the fraction of genes detected as allelically imbalanced at FDR = 0.001, stratified by dispersion level: low (θ < 0.005) and high (θ > 0.4); and gene expression levels– low (Mean(logTPM) < 3), medium (Mean(logTPM) ≥ 3 & Mean(logTPM) < 8) and high (Mean(logTPM) ≥ 8). **(C)** Performance comparison between ASPEN and scDALI using simulated data. Line plots illustrate TPRs calculated at FDR = 0.001 for genes categorized by gene expression and allelic dispersion levels (low dispersion – θ < 0.005, medium – θ ≥ 0.005 & θ < 0.4, and high – θ ≥ 0.4) across groups with varying degrees of deviation from the mean AR = 0.5. Only datasets simulated with a mean AR of 0.5 ± 0.2 are displayed. No performance differences were identified for groups with deviations from balanced allelic expression > 0.2. **(D)** Comparing performance in detecting allelic imbalance. TPRs are calculated based on the results from the simulated data with mean AR = 0.4. **(E)** Examples of the allelic ratio distributions of X-linked genes (*Tmsb4x*) and imprinted genes (*Meg3*). **(F)** Correlation of ASPEN-log_10_FDR values generated by running ASPEN mean test on the single-cell counts (x-axis) and pseudobulked counts (y-axis). Analyses were conducted in mouse cortical neurons (n = 972) from female Bl6 × Cast F1 hybrids and in CD8^+^ T cells at day 40 (n = 768) following LCMV infection in male Bl6 × Spret F1 hybrids.

To validate this framework, we performed extensive simulations across a range of gene expression levels, overdispersion parameters, and allelic levels. For realistic simulations, parameters of the beta-binomial model were estimated from a mouse brain organoid dataset ^34^.

ASPEN consistently demonstrated high sensitivity for detecting allelic imbalance with low false positives (**Fig. 3B, Supp. Table 1**). For low-dispersion genes, exemption from shrinkage improved sensitivity by up to 22%. In contrast, genes with high allelic dispersion benefited from shrinkage, which enhanced detection accuracy by up to 30%. We benchmarked ASPEN against scDALI ^24^, a method that models ASE via generalized linear mixed models, and found that ASPEN detected up to 30% more imbalanced genes across simulated datasets, with no detectable increase in false positives among allelically balanced controls (**Fig. 3C, Supp. Fig. 3A**). We also compared ASPEN’s method to the binomial test. As expected, as the binomial test does not account for overdispersion, the binomial test showed extremely high false positive rates in single cell droplet-based data (**Fig. 3D**). ASPEN runtimes for different gene and cell numbers are found in **Supp. Fig. 3B**.

We further assessed ASPEN’s ability to accurately resolve allele-specific expression using monoallelically expressed genes. In brain organoids derived from male *C57BL/6J × SPRET/Ei* F1 hybrids, ASPEN reliably identified 100% of expressed X-linked genes (43/43) as monoallelic. Additionally, all known imprinted loci, including *Meg3* and *Peg3*, were correctly recovered (n=10; **Fig. 3E, Supp. Table 2**), consistent with the ability to accurately resolve allele-specific expression.

We compared results between single-cell and bulk-level expression data. We aggregated scRNA-seq data into pseudobulk profiles across three major brain organoid subpopulations. ASPEN’s single-cell allelic imbalance calls showed strong concordance with pseudobulk-derived results, with 74–82% overlap in brain organoids and 77–93% agreement in T cells across different activation states (**Fig. 3F, Supp. Fig. 3C**). The slightly lower concordance in organoids may reflect a higher degree of cell-to-cell heterogeneity in allelic expression. This is consistent with prior observations of increased random monoallelic expression during embryonic stem cell differentiation ^39^ (**Supp. Fig. 3D**).

We applied ASPEN to distinguish allelic imbalance in T cells from *C57BL*/*6J × SPRET/Ei* F1 hybrids ^32^, identifying significant imbalance in 491 out of 809 expressed genes, including canonical markers of T-cell activation (*Cd69*), effector differentiation (*Gzmb, Ifng*), and memory formation (*Tcf7, Il7r*) (FDR < 0.05, **Supp. Table 3, Supp. Fig. 3E**). To explore potential regulatory mechanisms, we assessed whether allelic bias was associated with transcription factor (TF) motif differences. We used scATAC data from T-cells on day 7 post-LCMV infection from the *C57BL*/*6J* parental strain^32^, focusing on genes with strong allelic bias (ASPEN-mean FDR < 0.05, |log2FC| ≥ 1). For each parental haplotype, we calculated motif enrichment scores in promoter regions, comparing log odds scores for motifs significant on both alleles (*p* < 1×10^-4^). Across 53 genes and 312 motifs, we observed a mild correlation (ρ=0.2) between allelic bias and allele-specific TF motif enrichment (**Supp Fig 3F**). Integrating chromatin data from F1 hybrids would enable more direct mechanistic inference.

In summary, ASPEN showed strong concordance between single-cell and bulk-level allelic imbalance calls and recovered expected monoallelic and imprinted genes. It outperformed alternative methods in sensitivity and false positive control.

### Beyond imbalance: Modeling allelic variance

A key feature of ASPEN is its statistical framework to test for allelic over- and under-dispersion, enabling the identification of loci where allelic variance deviates from expression-level expectations (**Fig. 4A, Supp Fig 4, Methods**). Reduced levels of allelic variance may reflect transcriptional precision, while increased variance can signal regulatory plasticity, stochastic gene expression, or context-dependent modulation of cis-regulatory inputs ^35^.

**Figure 4.**
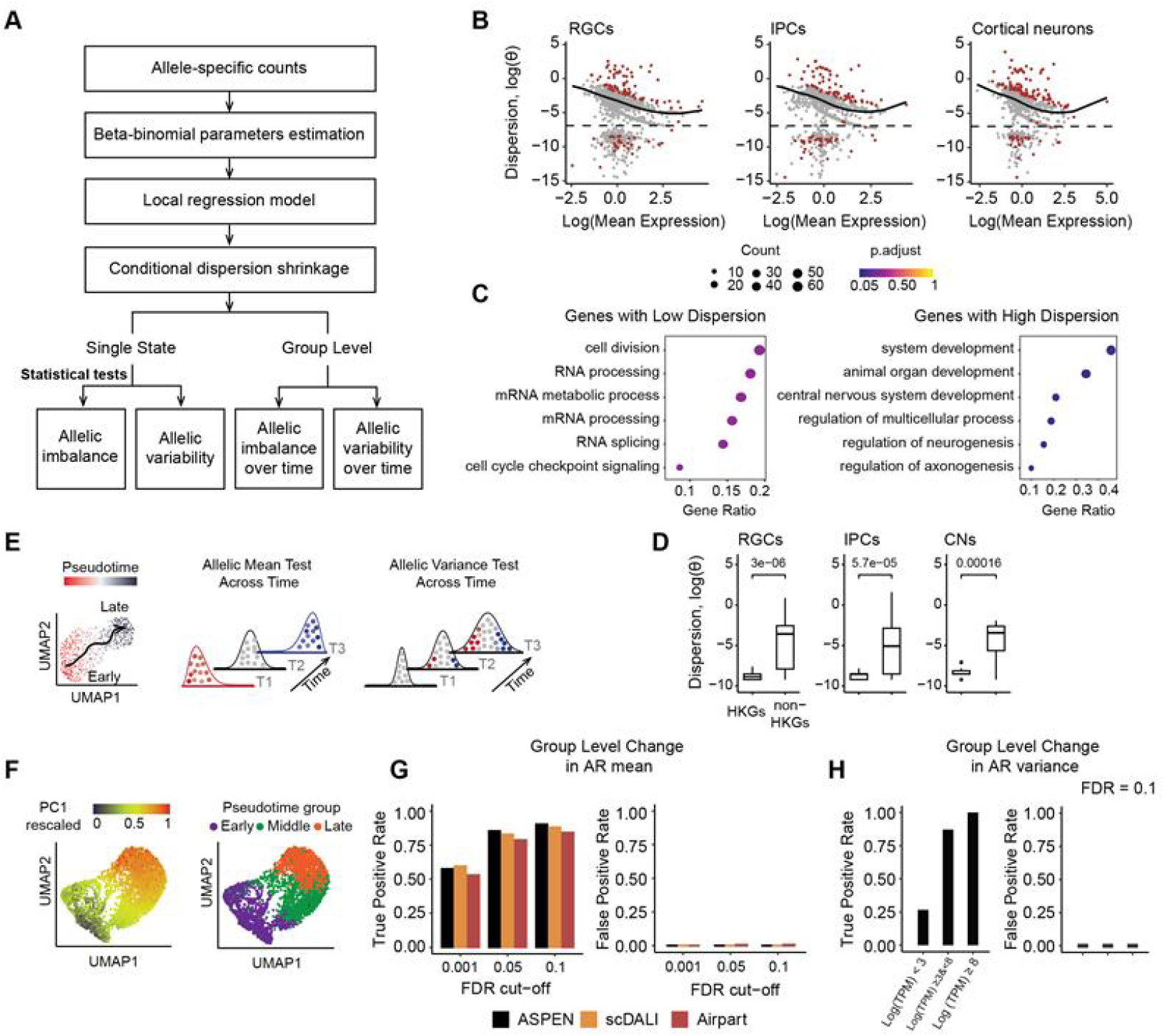
Detect allelic variation and differential changes in ASE with ASPEN. **(A)** Overview of the ASPEN analytical workflow. The main analysis steps involve the initial estimation of the allelic count distribution parameters, modeling allelic count dispersion as a function of gene expression, and conditional shrinkage toward the common dispersion estimates. **(B)** Results of the ASPEN var test in brain organoids data. Genes with significant deviation from expected variation levels are in red (FDR < 0.05): cortical neurons n = 140 (out of 973), IPCs n = 117 (out of 857) and RGCs n = 108 (out of 1,078). Genes below the dashed line were excluded from the trend modelling. **(C)** Top six Gene Ontology (GO) functional terms enriched in low dispersed genes (below the trend line) and highly dispersed genes (above the trend line). **(D)** Genes with housekeeping function in ESCs are lowly dispersion in different brain organoids cell types in female Bl6 × Cast F1 hybrids (RGCs: n = 26, IPCs: n = 20, CNs: n = 20, two-sided Wilcoxon rank-sum test). **(E)** Schematic representation of the ASPEN strategy used to detect changes in allelic mean or variance over time.**(F)** UMAP visualization of the simulated dataset used for testing. The cells are colored based on the rescaled PC1 coordinates representing temporal ordering (left) and by the cell assignments to discrete pseudotime bins: early, middle, and late (created by dividing the PC1 vector into tertiles) (right). **(G)** *Group-mean* test performance metrics. Test is applied to data simulated with group changes in allelic mean (0.45, 0.5, 0.55) vs. control (0.5, 0.5, 0.5). **(H)** G*roup-var* test performance metrics across different gene expression thresholds: low – Mean(logTPM) < 3, medium -Mean(logTPM)≥ 3 & < 8, high - Mean(logTPM) > 8. FDR = 0.1 was used for the evaluation. Data is simulated with changes in allelic variance (1×10^-3^, 0.05, 0.2) vs. control (1×10^-3^, 1×10^-3^, 1×10^-3^).

To detect such deviations, ASPEN first models the expected allelic dispersion as a function of gene expression, capturing the transcriptome-wide trend. It then evaluates whether individual genes significantly depart from this trend using an empirical permutation-based procedure applied to stabilized dispersion estimates (**Methods**). In female *C57BL*/*6J × CAST/Ei* F1 hybrid brain organoids, we identified 232 out of 1,261 expressed genes whose allelic variance significantly deviated from the expectation (**Fig. 4B**). Of these, 35% (83/232) were significantly under dispersed, while 65% (149/232) displayed significant over dispersion.

Genes with reduced allelic variance were enriched for core cellular processes, including RNA splicing, mRNA metabolism, and cell cycle regulation (FDR < 0.5; **Fig. 4C**, left panel), consistent with prior observations. Many have reported housekeeping function in ESCs ^36^ suggesting that allelic stability is selectively maintained in genes essential for cell function (Fisher’s exact OR = 1.42, *p* = 0.02, **Fig. 4D, Supp. Fig. 5A**). By contrast, over dispersed genes were enriched for neurodevelopmental pathways, including glial differentiation (*Sox4, Olig1, Ptn, Nfib*), axonogenesis (*Dcx, Pac3, Map1b, Nefm*), neurogenesis (*Vim, Nrp1, Sox11, Syt4*), and CNS development (*Tubb2a, Cenpf, Fabp7, Neurog2*) (FDR < 0.05; **Fig. 4C**, right panel).

To capture dynamic regulatory shifts during differentiation, ASPEN implements two likelihood ratio tests: one to detect temporal changes in allelic mean (*group-mean*) and another to test for temporal shifts in allelic variance (*group-var*) (**Fig. 4A, E**). In both cases, a null model assumes a shared set of beta-binomial parameters across timepoints, whereas the alternative model allows group-specific parameters. For the *group-mean* test, the mean is allowed to vary across the groups to capture distribution changes, while in *group-var*, the mean is fixed to assess differential changes in variance (**Fig. 4E**).

To benchmark the performance of ASPEN for group analyses, we simulated single-cell data along a pseudo-time axis (**Methods, Fig. 4F**). We divided the pseudotime vector into discrete equal-sized bins and cells were grouped into early, intermediate, and late stages. Simulations modelled: 1) group-level changes in allelic mean, 2) group-level changes in allelic variance, and 3) a null model with stable parameters (**Supp. Fig. 5B**).

ASPEN *group-mean* outperformed scDALI-Het and AirPart in detecting mean ASE differences with comparable specificity (sensitivity: 90%, 88%, 84%, respectively; specificity: 100%; FDR=0.1(**Fig. 4, Supp. Fig. 5C, Supp. Table 4**). ASPEN’s *group-var* test yielded 64% sensitivity and 100% specificity at FDR = 0.1 across all genes but increasing to > 87% sensitivity for more highly expressed genes (∼60% of genes) (**Fig. 4H, Supp. Fig. 5D, Supp. Table 5**).

Applying ASPEN to early neurogenesis in female *C57BL/6J × CAST/Ei* F1 hybrids revealed 162 genes with significant changes in mean allelic expression and 306 with differential variance, independent of total expression changes (**Supp. Fig. 5E; Supp. Table 6-7**). These included regulatory kinases (*Csnk1a1*), chromatin and transcriptional modulators (*Pbx1, Hmgb1, Fos*), and transcriptional co-regulators (*Taf1d, Ccna, Trim28r1, Eid1*). Notably, early progenitor populations exhibited the lowest variance (θ = 0.07 ± 0.01), which marginally increased in intermediate progenitors (θ = 0.09 ± 0.01) and differentiated cortical neurons (θ = 0.10 ± 0.02). This pattern was consistent with a reported increased in cellular heterogeneity during lineage commitment ^37^.

Eight genes with differential variance, including *Ankrd11, Eif3g, Nrxn1*, were high-confidence autism risk genes (SFARI database) ^38^. Additionally, five genes exhibiting variable allelic expression (*Eif3h, Lars, Oaz1, Selenok, Hsp90aa1*) have been implicated in anatomical brain phenotypes based on mouse knockout models ^39^ (**Supp. Fig. 5F**). We speculate that the temporal modulation of variance could reflect regulatory mechanisms that modulates gene expression programs during development and could be perturbed in disease states.

To summarize, ASPEN tests whether allelic variance at each gene departs from the transcriptome-wide expectation. This revealed a stabilization of variance in genes essential for core functions, highlighting regulatory flexibility in neurodevelopmental and disease-linked loci during differentiation.

### Detecting random and incomplete X inactivation

To identify genes with monoallelic expression, we used ASPEN’s estimation of beta-binomial shape parameters α and β. These parameters represent the support for expression from each allele, with values below 1 indicating cells showing monoallelic expression (**Fig. 5A**). When both α and β are below 1, cells express either one allele or the other, producing a bimodal allelic distribution consistent with random monoallelic expression (RME), in which a single allele is stochastically expressed in individual cells ^40–43^ (**Fig. 5A**).

**Figure 5.**
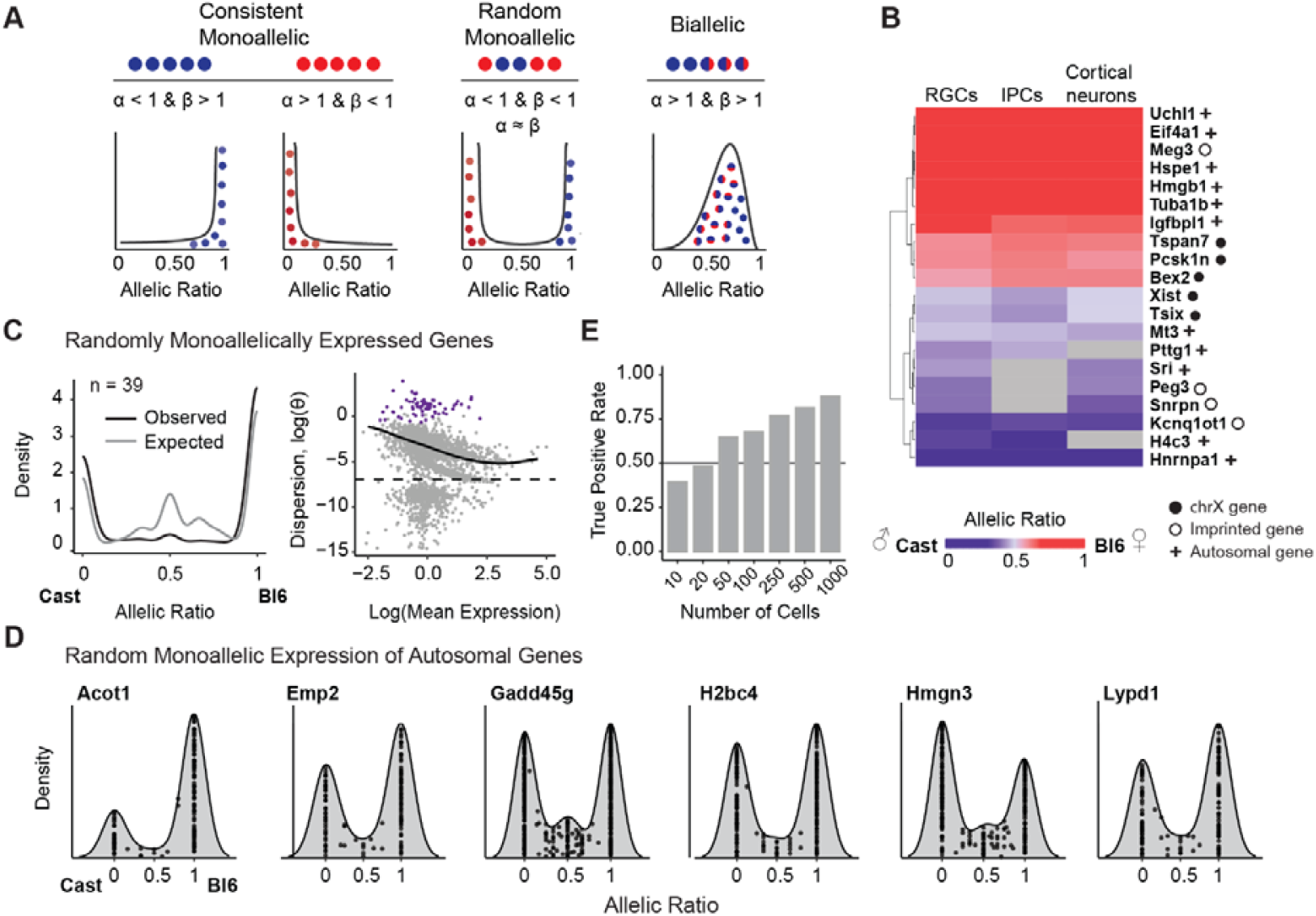
Monoallelic expression patterns in early neurogenesis. **(A)** Scenarios where allelic variance deviates from expected levels and the corresponding beta-binomial distribution shape parameters α and β. **(B)** The heatmap displays the mean allelic ratios across cells for genes with monoallelic expression. Genes were classified as monoallelic if their estimated beta-binomial α or β values were less than one. Results are from brain organoid data derived from female Bl6 × Cast hybrids using three cell types from the early neurodevelopmental pathway. Genes with allelic imbalance detected across all three cell types, with a mean expression above 1 in at least one, were selected for plotting (n = 18). Rows were hierarchically clustered using complete linkages. Genes without sufficient coverage in one of the cell types are indicated in grey. **(C)** Evaluating the expected versus observed dispersion distributions of genes with random monoallelic expression (RME) (total n = 39). The expected allelic ratio is calculated as the ratio of the simulated reference allele count to the total count. The expected gene set comprises genes without evidence of monoallelic expression, matched by expression to RME genes. The departure of the RME genes (in purple) from the common dispersion trend is visualized in the mean-variance plot. **(D)** Allelic ratio distribution plots for a selection of autosomal RME genes. **(E)** RME detection rates were low in the data simulated with a limited number of cells (n ≤ 20). RME simulation was performed with varying cell numbers (n = 10, 20, 50, 100, 250, 500, 1000).

Among 1,385 genes expressed in neuronal cells from female *C57BL/6J × CAST/Ei* F1 hybrids, 72 demonstrated monoallelic patterns, defined by extreme skew in the beta-binomial shape parameters, wherein either α or β was less than 1 (**Fig. 5A-B, Supp. Table 7**). Filtering based on these parameters identified canonical imprinted genes, including *Meg3, Peg3, Kcnq1ot1, Rian*, and *Snrpn* validating the approach (**Fig. 5**,**; Supp. Fig. 6A**).

To identify genes showing random monoallelic expression, we required both α and β < 1, with a difference between the parameters of less than 0.5 and further confirmed these candidates using the ASPEN variance test. We identified 39 high-confidence RME genes (**Fig. 5C**). Of these, 27 were X-linked, consistent with random X-inactivation dynamics in females. Five genes (*Bex2, Ndufb11, Pcsk1n, Sh3bgrl, Uba1*) were predominantly monoallelic in expression, yet they possessed a small proportion of cells (15-33%) displaying biallelic expression, consistent with incomplete X inactivation (**Supp. Fig. 6B**). Twelve RME genes were autosomal (**Fig. 5D; Supp. Fig. 6C**). These included *Gadd45g* and *H2bc4*, which we suggest are newly identified RME genes.

We examined whether expression level and cell numbers could have biased our results. RME genes were not more lowly expressed than non-RME genes (D = 0.182, p = 0.08, two-sided K-S test) but they were detected in fewer cells. To assess the impact of cell number level on ASPEN results, we simulated RME genes with different cell numbers, matching the distribution observed in real data. ASPEN achieved a sensitivity of 81.3% in simulations with a comparable number of cells (n=500) to the median cell numbers for RME genes detected by ASPEN in real data (n=526) (**Fig. 5E, Supp. Fig. 6D, 6E**). Notably, most autosomal RME genes recovered by ASPEN have been independently validated by clonal expansion followed by bulk RNA-seq, demonstrating ASPEN’s ability to detect RME *in vivo* (**Supp. Fig. 6F**) ^37,44,45^.

### ASPEN detects increased cis-regulatory control at housekeeping genes and key regulators of effector fate

Finally, we applied ASPEN to dissect the allelic regulatory dynamics during T-cell activation in acute viral infection, during which naïve CD8+ T cells transition through effector and memory states. Among 809 expressed genes, ASPEN identified 90 loci with allelic variance that deviated significantly from expression-level expectations (FDR < 0.05, **Fig. 6A**).

**Figure 6.**
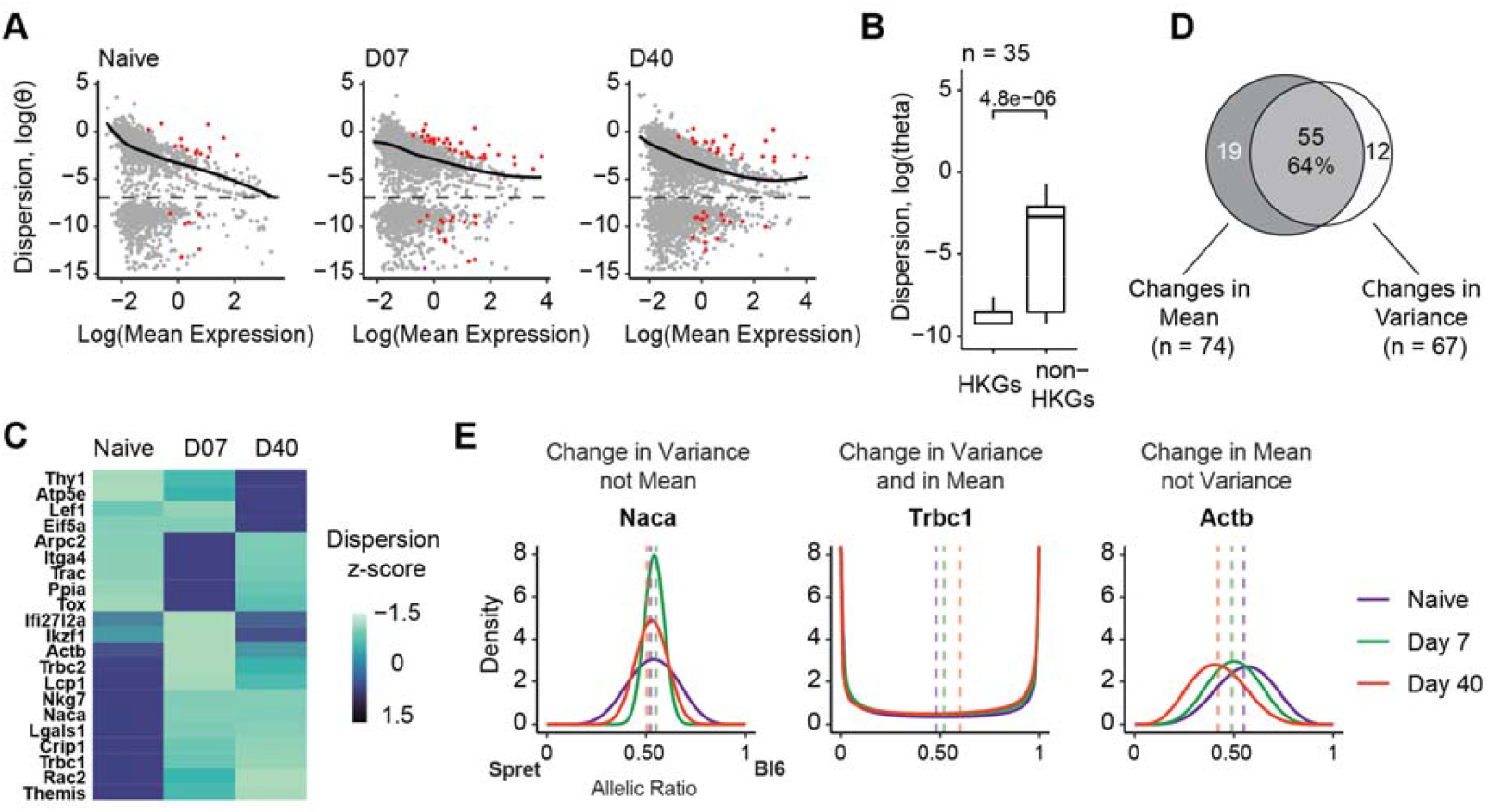
Evaluating allelic changes during T-cell activation. (**A**) Genes with significant deviations from the expected dispersion levels (in red) determined by the ASPEN var test (FDR < 0.05): Naïve n = 21, day 7 n = 50, day 40 n = 41. **(B)** Genes with housekeeping function had persistently low dispersion compared to background, matched by gene expression and number of detected cells (two-sided Wilcoxon rank-sum test). **(C)** Heatmap of the dispersion z-scores for the genes with ASPEN FDR var < 0.05 in at least one cell state (n = 21). Z-scores represent the empirical dispersion estimates for each gene scaled by column. **(D)** Venn diagram showing the overlap between the genes with dynamic changes in mean (ASPEN group-mean: FDR < 0.05) and variance (ASPEN group-var: FDR < 0.05). **(E)** Examples of differential changes in allelic distribution during T-cell activation. ASPEN tests categorize genes into three groups: those changing only in variance and not in mean (*Naca*, FDR group-var = 3.27 × 10^-8^, FDR group-mean = 1); those with changes in both mean AR and variance (*Trbc1*, FDR group-var = 2.57 × 10^-3^, FDR group-mean < 1.4 × 10^-96^); and those changing only in mean and not in variance (*Actb*, FDR group-var = 1, FDR group-mean < 1.4 × 10^-96^).

Consistent with observations in brain organoids, under-dispersed genes in T cells were also enriched for broadly conserved cellular processes (**Supp. Fig. 7A**), including housekeeping function (OR = 2.33, *p* = 2 × 10□^3^, Fisher’s exact test) (**Fig. 6B, Supp. Fig. 7B**). Furthermore, genes critical for CD8+ T cell clonal expansion and effector differentiation, such as *Il2ra* and *Cd8a*, exhibited significantly lower allelic variance relative to non-critical genes at matched expression level (Wilcoxon Z = –5.54, *p* = 3.04 × 10□□; **Supp. Fig. 7C**)^46^. In contrast, over dispersed genes were associated with immune-specific functions. These included pathways related to immune system processes (*Trac, Tox, Nkg7, Ccl5*), lymphocyte activation (*Lgals1, Actb, Themis, Lef1*), and leukocyte signaling (*Myo1f, Klrd1, Thy1, Ccnd3*) (**Supp. Fig. 7A**).

Using ASPEN’s group-var test, we identified a shift in allelic variance over the course of T-cell activation. Naïve cells exhibited the highest allelic variance (θ = 0.12 ± 0.04), which progressively declined at day 7 post-infection (θ = 0.05 ± 0.01) and further decreased in memory cells at day 40 (θ = 0.02 ± 0.01) (**Fig. 6C, Supp. Fig. 7D**). Among 119 differentially expressed genes, ASPEN identified significant temporal changes in allelic mean ratios for 74 genes (62%) and differences in allelic variance for 67 genes (56%), with 55 genes showing coordinated changes in both parameters (FDR < 0.05, **Fig. 6D, Supp. Table 9**).

Genes exhibiting differential allelic regulation showed different patterns of allelic expression (**Fig. 6E, Supp. Fig. 7E**). For example, *Naca*, nascent polypeptide-associated complex subunit alpha involved in T cells proliferation, showed changes in variance but not in allelic ratio (**Fig. 6E**). In contrast, *Actb* demonstrated a shift in allelic preference without concomitant changes in variance, while *Trbc1* transitioned from biallelic to maternally biased expression while maintaining stable total transcript levels (**Fig. 6E**).

Together, these examples highlight the diversity of expression dynamics. Allelic variance decreased as cells were activated, suggestive of a tightening of transcriptional control with functional specialization. Core effector genes for T cell activation exhibited low variances independent of expression level suggestive of regulatory control.

## Conclusion

Understanding heterogeneity in cis-regulatory control is central to decoding how gene expression varies within and across cell states. ASPEN provides a general framework to uncover these dynamics at single-cell resolution from F1 hybrids. By integrating a sensitive mapping pipeline with moderated beta-binomial modeling and adaptive shrinkage, ASPEN improves the detection of both allelic imbalance and variance, outperforming existing methods in sensitivity and false discovery control.

Beyond identifying shifts in mean allelic ratios, ASPEN quantifies allelic variance. It detects reduced variance in housekeeping and essential genes, suggesting tight cis-regulatory control, and increased variance in neurodevelopmental and immune-related genes, indicative of stochastic or context-dependent regulation. ASPEN also enables the detection of dynamic changes in allelic regulation across differentiation and activation states, uncovering genes with random monoallelic expression and incomplete X inactivation. These insights extend to rare or postmitotic cell types, where traditional clonal expansion approaches are infeasible. These findings demonstrate allelic variance is a key dimension of cis-regulatory control, and that ASPEN provides a set of robust methods for uncovering these regulatory programs.

## Methods

We generated F1 alignment indices for four common mouse hybrid strains: *C57BL/6J* (Bl6) *× Mus Castaneus (CAST/Ei)* (Cast), Bl6 × *Mus musculus molossinus (MOLF/Ei)* (Molf), Bl6 × *Mus musculus musculus (PWK/Ph)* (Pwk), and *C57BL*/*6J × Mus Spretus (SPRET/Ei)* (Spret). Reads from hybrid animals were aligned to their parental combined genomes. Only reads that could be allelically resolved were used in gene expression quantitation.

### Combined genome generation

We generated combined genomes containing two sets of chromosomes for each genetic background, which allowed for unambiguous assignment of allelic reads. Variant call files (release 5) for four wild-derived mouse strains (*CAST/Ei, MOLF/Ei, PWK/Ph*, and *SPRET/Ei*) were downloaded from the Mouse Genome Project ^47^. Strain-specific genomes were created using high-quality calls (filter set to PASS, GT = “1/1”) and incorporating (i) only SNPs and (ii) both SNPs and insertions and deletions (indels) into the reference *C57BL/6J* genome (GRCm38, release 68) with BCFtools consensus (v1.12) ^48^. Each genome was concatenated with the reference *C57BL/6J* genome to produce combined genomes for each mouse strain. The chain files from the BCFtools were used to create strain-specific gene annotations by converting *C57BL/6J* gene annotations (Ensembl release 102) via UCSC liftOver ^49^. Combined genome index files were generated using STAR (v2.7.6a) ^50^.

### Beta-binomial modeling of allelic counts

To quantify allele-specific expression, ASPEN requires as input two matrices for each cell and gene: the number of reads mapping to the reference allele (m_j_ and the total read count (n_j_), where i indexes genes and j indexes cells. For each gene, ASE is modeled using a beta-binomial distribution, parameterized by a mean allelic ratio (µ_i_) and a dispersion parameter (θ_i_), which together capture the probability and variability of observing reference allele counts across cells. The likelihood of observing m_j_ reference reads out of n_j_ total reads for gene *i* in cell *j* is given by:

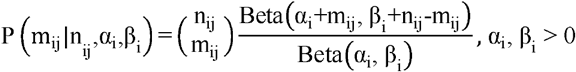

where α_i_ and β_i_ are the shape parameters of the beta prior. These are related to the mean and dispersion by:

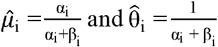

Here, 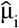 represents the average allelic ratio for gene *i*. and 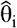 reflects the overdispersion or variability in allelic ratios across cells beyond binomial sampling noise ^51^.

Parameter estimation is performed by maximizing the likelihood function:

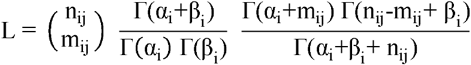

Maximum likelihood estimation (MLE) was performed with R (v4.3.1)^52^ ‘optim’ function using BFGS quasi-Newton method with positive constraints on α_i_ and β_i_.

### Modeling the relationship between allelic dispersion and mean expression

ASPEN moderates gene-wise dispersion estimates by leveraging the overall relationship between dispersion and mean expression across all genes to address sampling noise and technical variation. This information-sharing uses a weighted log-likelihood framework^33,34,48^:

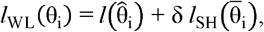

where 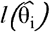 is the log-likelihood for the gene-specific dispersion estimate, 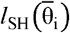 is the log-likelihood for the shared dispersion at matched expression level, and δ determines the relative influence of the shared trend.

To estimate the shared dispersion values, we model the dependence of gene-level dispersion, 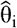,as a function of gene expression (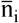 the average total read count per gene) using local regression implemented in the locfit (v.1.5-9.8^53^) R package:

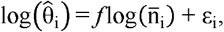

where *f* captures the smoothed mean-variance relationship and ε_i_ is the residual for gene *i*. All genes are included in the regression to define a representative mean-variance trend. The resulting fitted values 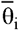, represent the expected dispersion for genes expressed at similar levels.

### Bayesian shrinkage implementation

ASPEN applies empirical Bayes shrinkage to stabilize dispersion estimates by combining gene-specific information with the transcriptome-wide mean-variance trend. For each gene, the posterior (shrunken) dispersion is calculated as a weighted average of the individual and local trend estimates, with the weight parameter (δ) controlling the degree of shrinkage. To obtain the posterior (shrunken) dispersion for gene *i*, we modified the analytical solution which was derived for the double binomial distribution and extended to the beta-binomial distribution ^48^:

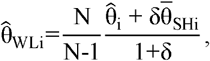

where 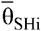 is the local trend, obtained by local regression on log-dispersion versus log-expression, serving as a prior for each gene’s dispersion, 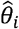 is the individual gene dispersion estimate, N is the effective degrees of freedom, and δ is weight given to the shared likelihood. 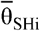 is treated as prior for θ_i_ under a Bayesian hierarchical framework. To avoid overfitting, the shrinkage weight is adjusted by the factor N-K, where K is the number of fitted parameters, yielding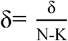.

Hyperparameters for shrinkage are determined by fitting a gamma distribution to the observed dispersions across genes using maximum likelihood. All optimization is performed in R (v4.3.1) using the BFGS method ^52^. Initial parameters were set to N = 20 and δ = 10. For all real and simulated data, we set δ = 50 and N = 30, as determined empirically from T cell data.

### Treatment of genes with low allelic variation

We applied selective shrinkage for genes with low dispersion estimates that pass the read count threshold (≥ 5 cells ≥5 UMIs) which are moderated towards the common trend. To detect these lowly variable genes, we plot estimated dispersion against the mean expression. We fit a smoothing function across all data points using the local regression residuals, ε_i_:

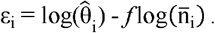

Low variance genes are the outliers that cluster distinctly from the model trend. To provide a more principled approach, we also applied the median absolute deviation-squared, MAD^2^ statistic ^54^,

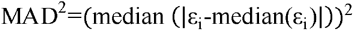

and defined lowly variable genes as those with ε_i_ < MAD^2^. Classification of lowly variable genes used the threshold of θ < 0.001 for the real data and θ < 0.005 for the simulated data.

We note that for allelic tests, the dispersion trend was modelled after excluding these lowly variable genes from Bayesian shrinkage and updating the shared estimates, 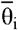.

### Statistical framework for testing allelic imbalance

After estimating gene-specific beta-binomial parameters and obtaining shrunken dispersion values, ASPEN tests for allelic imbalance using a log-likelihood ratio test. For each gene, the null hypothesis (H_0_) assumes allelic ratios follow a beta-binomial distribution with a fixed mean (*x*), reflecting the expected background allelic ratio, and the estimated dispersion. The alternative hypothesis (H_1_) allows the mean allelic ratio 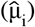 to be estimated from the data:

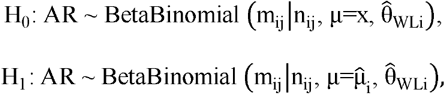

where x is the fixed allelic ratio which is determined based on empirical background (except for simulated data where x = 0.5, representing theoretical allelic balance), 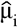 is the allelic ratio and 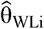 is the shrunken dispersion that are specific to each gene. Both null and alternative models are fit by maximum likelihood in R as above.

For low-variance genes, the empirical (unshrunken) dispersion estimate, 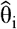 is used instead of the shrunken value 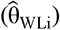.The fixed allelic ratio x is set to the global mean across all autosomal, non-imprinted genes passing quality filters, or to 0.5 in simulated data. A likelihood ratio test (LRT) is then performed, comparing the fit of the null and alternative models. Test statistics are evaluated using the χ^2^ distribution with one degree of freedom, and p-values are corrected for multiple testing using the false discovery rate (FDR) ^55^.

### Statistical framework for testing changes in allelic variance

For each gene, we quantified allelic variation in two steps. First, we establish the baseline level of expected variation 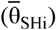 using local regression to model the relationship between expression level and allelic variation across all genes. We then compared each gene’s observed stabilized dispersion 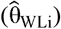 against this, holding the mean allelic ratio 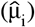 constant. Both dispersion and mean parameters were estimated by MLE. By analyzing the fit of these models, the test distinguishes genes where ASE patterns vary between cells due to biological processes rather than technical noise.

To detect genes with significantly greater or lesser allelic variance than expected for their expression level, we performed a likelihood ratio test based on the following hypotheses:

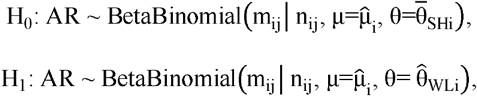

where 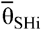 is the predicted baseline dispersion and 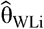 is the gene’s stabilized (shrunken) dispersion. Shrinkage is applied to all genes meeting minimum filters (≥ 5 cells ≥5 UMIs). For each gene, we used a permutation-based approach to estimate empirical p-values: reference allele counts were simulated under the null model, and the observed likelihood ratio was compared to the distribution of simulated ratios. We found that 500 permutations were sufficient for stable p-value estimates (tolerance <0.001). *p*-values were adjusted using the FDR ^55^. We noted that the simulation-based *p*-value converged around 400 permutations with a tolerance of 0.001 (**Supp. Fig. 8**). Hence, we ran 500 permutations.

### Detecting group-level changes

ASPEN implements two LRTs to assess temporal allelic changes: group-mean (for changes in mean allelic ratios) and group-var (for changes in allelic variance) across discrete groups. Both tests require cells to be assigned to discrete groups, such as by cluster, cell type, or discretized pseudotime. Beta-binomial parameter estimation, dispersion modeling, and shrinkage are performed at both global (all cells) and group-specific levels for each gene.

#### Group-mean test

To assess changes in mean allelic ratio while controlling for dispersion, the null hypothesis assumes all groups share the same mean and shared (‘global’) dispersion:

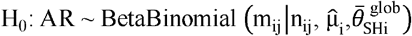

where 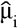 denotes the shared-group mean and where 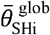 is the shared dispersion. The alternative hypothesis allows the mean to vary between groups and dispersion is matched to expression based on shrinkage:

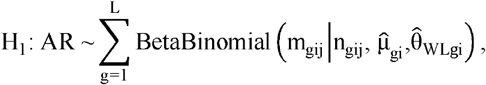

where 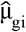 is the group-specific mean and 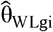 the group-specific shrunken dispersion estimate for gene i.

#### Group-var test

To assess changes in allelic variance while holding the mean constant, the null hypothesis assumes all groups share the same mean and dispersion:

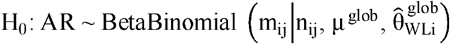

where μ^glob^ is the global mean allelic ratio. The alternative hypothesis allows the dispersion parameter to vary between groups:

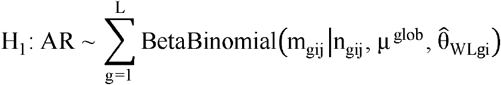

is 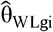 the group-specific shrunken dispersion estimate.

For both tests, maximum likelihood estimation was used to fit the model parameters, and statistical significance was determined by a likelihood ratio test with *p*-values evaluated from the 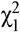 distribution (1 degree of freedom). Multiple testing correction is performed using the Benjamini-Hochberg FDR procedure ^55^.

### Generation of simulated data

To simulate single-cell RNA-seq data, we estimated model parameters using the brain organoid dataset from male *C57BL/6J* × *SPRET/Ei* F1 hybrids (Bl6 × Spret) ^34^. This dataset included 3,269 cells (clone four), spanning three cell types. The top 2,000 most variable genes identified using getTopHVGs from scater (v1.28.0) ^56^, and parameters were estimated by fitting a zero-inflated negative binomial model (ZINB-WaVE model, v.1.22.0) ^57^ to the total count matrix, specifying K□=□2 latent cell-level covariates and using default values for other settings.

Reference allele counts were simulated using VGAM (v.1.1-9) ^58^ by drawing beta-binomial counts from the simulated total count matrix. To assess ASPEN’s ability to detect genes with allelic imbalance, we generated multiple datasets with allelic ratios set to {0.10, 0.30, 0.35, 0.40, 0.42, 0.45, 0.46, 0.47, 0.48, 0.49, 0.50, 0.51, 0.52, 0.53, 0.54, 0.55, 0.57, 0.60, 0.65, 0.70, 0.90}. The dispersion estimates from real data were kept fixed for the respective gene in each simulation. The simulation for each group was repeated five times to improve technical reproducibility.

To simulate temporal or pseudotime scenarios, we again used the ZINB-WaVE total count matrix. Cells were processed using Seurat (v. 4.0.5) ^59^. Counts were log-normalized and scaled to 10,000 counts, followed by PCA on all genes. The top 20 principal components were used for Louvain clustering (FindClusters), and UMAP was performed for visualization. Pseudotime was defined by rescaling PC1 coordinates between 0 and 1, then dividing cells into three equal-sized bins (tertiles).

To test detection of temporal changes, beta-binomial counts were simulated for three bins. In the first scenario, the mean allelic ratios were set to {0.45, 0.5, 0.55} with constant dispersion {1×10^-3^, 1×10^-3^, 1×10^-3^} across bins. In the second scenario, mean ratios were held constant (0.5) while dispersion varied across bins {1×10^-3^, 0.05, 0.2}. In control simulations, both mean and dispersion were fixed (0.5, 1×10^-3^) in all bins.

For assessing the robustness of random monoallelic expression (RME) detection, we sampled total counts from real RME genes and subsampled columns to create datasets with varying cell numbers (n = 10, 20, 50, 100, 250, 500, 1000). Each subsampling was repeated 30 times, with a minimum of 500 genes per set. To mimic RME, allelic counts were sampled using rbetabinom.ab in VGAM (v.1.1-0)^58^ with shape parameters (alpha, beta) obtained from ASPEN estimates for real RME genes and limit.prob set to 0.5.

### Detecting allelic imbalance in the simulated data

We applied scDALI (v0.1.0) ^24^ in homogeneous (Hom) mode to the simulated counts to evaluate the global allelic imbalance. In ASPEN, we used the mean test following the dispersion modeling and shrinking steps. For both methods, we tested the observed allelic ratio against the null hypothesis of AR = 0.5. *P*-values were adjusted for multiple testing via the false discovery rate (FDR). To assess the performance of the tests, we stratified genes by their expression levels based on transcript per million (TPM) values and allelic dispersion. The top 10% of genes fell into the high expression group (log(TPM) > 8), the next 50% to the medium expression group (log(TPM) < 8), and the remaining genes were in the low expression group (log(TPM) < 3). The TPM values were obtained by running calculateTPM from scuttle (v1.10.2) ^56^ on the total count matrices. Genes were partitioned based on allelic dispersion as follows: genes whose dispersion was less than 0.005 were assigned to the low-dispersion group, and a threshold of 0.45 was selected to assign genes to either the medium- or high-dispersion group. For differential allelic expression benchmarking (*group-mean*), scDALI was used in heterogeneous (Het) mode and allelicRatio function was used from Airpart^23^ with DAItest set to TRUE.

### Bulk RNA-Seq data processing

Mouse fibroblast RNA-Seq fastq files to evaluate read mapping strategies were downloaded from GSE193728. To compare the distribution of genes with low dispersion between single-cell and bulk RNA-seq data, CD8+ T-cell data from male Bl6 × Spret hybrid data were obtained from GSE126770. The reads were trimmed via Trimmomatic ^60^ in pair-end mode and mapped to the combined genome index via STAR (v2.7.6a) ^50^. To achieve full-length read alignment, the insertions, soft-end clipping and splicing modes were turned off (--outBAMcompression 6 -- outFilterMultimapNmax 1 --outFilterMatchNmin 30 --alignIntronMax 1 --alignEndsType EndToEnd --outSAMattributes NH HI NM MD AS nM --scoreDelOpen -1000 --scoreInsOpen -1000). Reads mismatched (flagged as ‘NM’) and reads overlapping diverse strain-specific regions that were enriched for recently transposed long interspersed nuclear elements (LINEs) and long-terminal repeat (LTR) elements^61^ were removed. Reads were counted via Rsubread (v2.4.3104)^62^ with the following parameters: isPairedEnd=T, allowMultiOverlap=T, and countMultiMappingReads=F. To normalize for sequencing depth, size factors were estimated via total counts (sum of allelic counts) with DESeq2 (v1.30.193) ^30^ and used to adjust allelic counts.

### scRNA data preprocessing

CD8+ T-cell data for naïve, day 7 and day 40 postinfection samples were obtained from the GSE164978 dataset (SRA files SRR13450127 and SRR13450128). Fastq files for mouse brain organoids generated from female Bl6 × Cast and male Bl6 × Spret F1 hybrids were downloaded from GSE268332. Files were mapped to the respective strain genomes via STARSolo (v.2.7.6a) ^63^ (--soloType CB_UMI_Simple --soloUMIlen 12 --soloCBstart 1 --soloCBlen 16 --soloUMIstart 17 --soloCBwhitelist <(zcat ${dir}/align/3 M3M-february-2018.txt.gz) --soloFeatures Gene -- outSAMattributes NH HI nM AS CR UR CB UB GX GN sS sQ sM NM MD -- outBAMcompression 6 --outFilterMultimapNmax 1 --outFilterMatchNmin 30 --alignIntronMin 20 --alignIntronMax 20000 --alignEndsType EndToEnd --outFilterMismatchNoverReadLmax 0.001 -- scoreDelOpen -1000 --scoreInsOpen -1000). To evaluate Expectation Maximization strategy for the multimapping reads allocation, we added the following flags in STARSolo: -- outFilterMultimapNmax 3 and --soloMultiMappers EM. The total count matrix was produced by combining maternal and paternal allelic counts. Genes that were expressed in fewer than 10 cells were removed from further analysis.

### Single cell data allelic testing in CD8+ T cells

We conducted ASPEN allelic imbalance and variance tests on data from F1 mouse brain organoids and CD8 T cells. We examined 5,095 genes across 723 (naïve), 400 (day 7), and 582 (day 40) cells. The total count matrices for each time point were merged. Size factors were calculated using computeSumFactors from scran (v.1.28.2) ^64^ by pooling similar groups of cells together (we used the original time points for cluster assignment). The same scaling was applied to both the total and reference allele counts. Initial beta-binomial parameters were estimated, and dispersion modeling and shrinkage were conducted with raw counts. The expected allelic ratio was adjusted to the empirical ratio of 0.52 for the allelic imbalance test to reflect an alignment bias toward the reference genome (**Supp Fig. 9A**). Testing was then performed on the normalized counts, selecting genes that had at least 5 cells with at least 5 mapped reads. We ordered the datasets by time points (naïve, day 7, and day 40 post-infection) to assess dynamic changes in the allelic ratio distribution and used this as a discrete temporal assignment. P values for the ASPEN tests were FDR-adjusted.

### Pseudobulked allelic testing in CD8+ T cells

Merged CD8+ T-cell counts were processed in Seurat (v. 4.0.5) ^59^. We performed principal component analysis (PCA) on log-normalized data and scaled it to 10000 counts using all genes. The top 20 principal components (PCs) were selected for downstream Louvain clustering as implemented in the FindClusters function (resolution = 0.9), and low-dimensional representations were obtained through uniform manifold approximation and projection (UMAP). This process produced three clusters per time point, which were then used to aggregate the total counts as pseudoreplicates within each cell state (in lieu of the biological replicates) (**Supp Fig. 9B**). The same cell assignment was used to pool the reference allele counts. Size factors were estimated via total counts with DESeq2 (v1.30.1) ^30^ and were applied to scale the total and reference allele counts in parallel. Genes with a minimum of 10 reads across all clusters within the sample were retained for allelic imbalance testing. We checked for mapping bias in the pseudobulk dataset, revealing a slight skew toward the Bl6 allele, which was accounted for by adjusting the expected ratio (**Supp Fig. 9C**). P-values for the allelic imbalance test obtained from ASPEN were FDR-adjusted.

### Analysis of mouse brain organoid single-cell RNA-seq data

To obtain initial beta-binomial parameters and shrunk dispersion values from CD8 T cells, we used the same procedure as for the male Bl6 × Spret hybrid data. For allelic imbalance testing, we adjusted the allelic ratio value under the null hypothesis to 0.54. Using beta-binomial shape parameters, we defined genes with α < 1 OR β < 1 as monoallelically expressed.

### Allelic imbalance in pseudobulked counts from mouse brain organoids

Single-cell Bl6 alleles and total counts from four female Bl6 × Cast F1 hybrids were aggregated by clonal origin to generate pseudobulk replicates. We estimated size factors using total counts with DESeq2 (v1.30.1) ^30^ and applied them to scale the total and Bl6 allele counts. Genes with fewer than 10 total counts across the pseudoreplicates were excluded from further analysis. Differentially expressed genes between RGC and IPC cell types were identified based on the total pseudobulked counts using nbinomWaldTest function from DESeq2 (v1.30.1) ^30^.

### Identification of random monoallelic expression

We identify RME expression patterns using the shape parameters (α and β) of the beta-binomial distribution. When either α or β falls below one, the distribution becomes skewed towards one allele. The hallmark of RME emerges when both α and β are less than one and are approximately equal, producing a U-shaped allelic density – indicating that individual cells express either the maternal or paternal allele but rarely both. We identified incomplete X chromosome inactivation in genes that exhibited biallelic expression based on the following criteria: mean allelic ratio between 0.25 and mean 0.75 in at least 15% of cells, with a minimum of five cells.

### Allele-specific TF binding motif enrichment analyses

Processed open chromatin data for T cells at day 7 post-LCMV infection were obtained from GSE164978^32^. scATAC peaks overlapping the promoter regions (+/-1 Kb) around TSS of the allelically imbalanced genes (ASPEN-mean FDR < 0.05 & |log2FC| ≥ 1) were selected. Coordinates of the corresponding regions on the Spret allele were obtained by lifting over the Bl6 promoter region coordinates using the chain file generated above and USCS liftOver ^49^. To calculate the motif enrichment scores, FIMO (v.5.5.4) ^65^ was run for Bl6 and Spret sequences of their respective promoter regions. The sequences were extracted from either the Bl6 or Spret genomes using the getfasta command from bedtools^66^. Selecting motif with significant enrichment on both alleles (*p* < 1 ×10^−4^), the differences between the FIMO log odds scores were calculated.

### Gene stability analyses

A list of genes ranked by the stability index was obtained from Lin et al^33^. Genes with Stability Index ≥ 0.8 were defined as Stably Expressed Genes (SEGs) and genes with Stability Index < 0.8 –as non-SEGs. We matched the SEGs and non-SEGs by their expression level and the number of available cells by performing the nearest neighbour search using MatchIt (v.4.5.5) ^67^ with parameters set to method = “nearest” and distance = “glm”. Housekeeping genes were obtained from HRT Atlas v1.0^36^ database by selecting either “Embryonic Stem Cells” or “Spleen bulk tissue”. Essential and non-essential genes for T-cell clonal expansion were selected in the same manner. The matching was performed based on the gene expression only.

### Functional enrichment analysis

Gene Ontology (GO) term enrichment analysis was performed in clusterProfiler (v4.8.3) ^68^ via the org.Mm.eg.db database (3.17.0) ^69^. We used the enrichGO function to identify enriched biological processes in genes with low dispersion 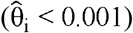 in the T-cell data collected on day 7 post-LCMV infection. All genes with sufficient coverage from the same time point were used as the background for GO enrichment.

## Availability of data and materials

ASPEN is available as an R package at https://github.com/ewonglab/ASPEN. Single-cell RNA-seq data for CD8 T cells from naïve, day 7, and day 40 LCMV Armstrong infection samples were downloaded from the GSE164978 dataset (SRA: SRR13450127, SRR13450128) ^32^. Bulk RNA-Seq data for CD8+ T cells 7 days after LCMV Armstrong infection were downloaded from GSE126770 ^31^. Fastq files for bulk RNA-seq of mouse fibroblasts from four F1 hybrids (Bl6 × Cast, Bl6 × Molf, Bl6 × Pwk, and Bl6 × Spret) were obtained from GSE193728 ^70^. Single-cell RNA-seq data for mouse brain organoids (Bl6 x Cast and Bl6 x Spret) were downloaded from GSE268332 ^34^. Variant call files (release 5) between *C57BL/6J* and four wild-derived mouse strains (*CAST/Ei, MOLF/Ei, PWK/Ph*, and *SPRET/Ei*) were obtained from the Mouse Genome Project ^47^. A list of recently transposed long interspersed nuclear elements (LINEs) and long-terminal repeat (LTR) elements in the strain-specific genomes was downloaded from the original publication ^61^.

## Acknowledgments

We thank Daniel Medina-Cano and Mohammed Islam for data used for method evaluation. This project was undertaken with assistance from the resources and services of the National Computational Infrastructure (NCI), supported by the Australian Government. We are grateful for the support of the UNSW NCI Resource Allocation Scheme (doi:10.26190/PMN5-7J50).

## Authors’ contributions

E.S.W. conceived and supervised the project. V.P. wrote the code except for the variance test and performed all methods evaluation. T.V. supplied data for method evaluation. E.S.W. wrote the manuscript with input from V. P. M.N. wrote the code for the variance test. All the authors reviewed the manuscript.

## References

1. Babak, T. et al. Global Survey of Genomic Imprinting by Transcriptome Sequencing. Current Biology 18, 1735–1741 (2008).

2. Goncalves, A. et al. Extensive compensatory cis-trans regulation in the evolution of mouse gene expression. Genome Res. 22, 2376–2384 (2012).

3. Andergassen, D., Smith, Z. D., Kretzmer, H., Rinn, J. L. & Meissner, A. Diverse epigenetic mechanisms maintain parental imprints within the embryonic and extraembryonic lineages. Developmental Cell 56, 2995-3005.e4 (2021).

4. Reinius, B. et al. Analysis of allelic expression patterns in clonal somatic cells by single-cell RNA–seq. Nat Genet 48, 1430–1435 (2016).

5. Keniry, A. et al. BAF complex-mediated chromatin relaxation is required for establishment of X chromosome inactivation. Nat Commun 13, 1658 (2022).

6. Coolon, J. D., McManus, C. J., Stevenson, K. R., Graveley, B. R. & Wittkopp, P. J. Tempo and mode of regulatory evolution in Drosophila. Genome Research 24, 797 (2014).

7. Khoueiry, P. et al. Uncoupling evolutionary changes in DNA sequence, transcription factor occupancy and enhancer activity. eLife 6, e28440 (2017).

8. Wong, E. S. et al. Interplay of cis and trans mechanisms driving transcription factor binding and gene expression evolution. Nature Communications (2017).

9. Hu, C. K. et al. cis-Regulatory changes in locomotor genes are associated with the evolution of burrowing behavior. Cell Reports 38, 110360 (2022).

10. Wittkopp, P. J., Haerum, B. K. & Clark, A. G. Evolutionary changes in cis and trans gene regulation. Nature 430, 85–88 (2004).

11. Wang, X., Werren, J. H. & Clark, A. G. Allele-Specific Transcriptome and Methylome Analysis Reveals Stable Inheritance and Cis-Regulation of DNA Methylation in Nasonia. PLoS Biology 14, e1002500 (2016).

12. Rossi, M. et al. Visual mate preference evolution during butterfly speciation is linked to neural processing genes. Nat Commun 11, 4763 (2020).

13. Springer, N. M. & Stupar, R. M. Allele-Specific Expression Patterns Reveal Biases and Embryo-Specific Parent-of-Origin Effects in Hybrid Maize. The Plant Cell 19, 2391 (2007).

14. Zhang, X. & Borevitz, J. O. Global Analysis of Allele-Specific Expression in Arabidopsis thaliana. Genetics 182, 943 (2009).

15. Bell, G. D. M., Kane, N. C., Rieseberg, L. H. & Adams, K. L. RNA-Seq Analysis of Allele-Specific Expression, Hybrid Effects, and Regulatory Divergence in Hybrids Compared with Their Parents from Natural Populations. Genome Biology and Evolution 5, 1309–1323 (2013).

16. Tirosh, I., Reikhav, S., Levy, A. A. & Barkai, N. A Yeast Hybrid Provides Insight into the Evolution of Gene Expression Regulation. Science 324, 659–662 (2009).

17. Bueker, B. et al. Meiotic recombination in the offspring of Microbotryum hybrids and its impact on pathogenicity. BMC Evolutionary Biology 20, 123 (2020).

18. Tsouris, A., Brach, G., Schacherer, J. & Hou, J. Non-additive genetic components contribute significantly to population-wide gene expression variation. Cell Genomics 4, 100459 (2024).

19. Jiang, Y., Zhang, N. R. & Li, M. SCALE: modeling allele-specific gene expression by single-cell RNA sequencing. Genome Biology 18, 74 (2017).

20. Larsson, A. J. M. et al. Genomic encoding of transcriptional burst kinetics. Nature 565, 251–254 (2019).

21. Pacini, G. et al. Integrated analysis of Xist upregulation and X-chromosome inactivation with single-cell and single-allele resolution | Nature Communications. Nature Communications 12, 3638 (2021).

22. Lentini, A. et al. Elastic dosage compensation by X-chromosome upregulation. Nat Commun 13, 1854 (2022).

23. Mu, W. et al. Airpart: interpretable statistical models for analyzing allelic imbalance in single-cell datasets. Bioinformatics 38, 2773–2780 (2022).

24. Heinen, T. et al. scDALI: modeling allelic heterogeneity in single cells reveals context-specific genetic regulation. Genome Biology 23, 8 (2022).

25. Qi, G. et al. Single-cell allele-specific expression analysis reveals dynamic and cell-type-specific regulatory effects. Nat Commun 14, 6317 (2023).

26. Saukkonen, A., Kilpinen, H. & Hodgkinson, A. Highly accurate quantification of allelic gene expression for population and disease genetics. Genome Res 32, 1565–1572 (2022).

27. Robinson, M. D. & Smyth, G. K. Moderated statistical tests for assessing differences in tag abundance. Bioinformatics 23, 2881–2887 (2007).

28. McCarthy, D. J., Chen, Y. & Smyth, G. K. Differential expression analysis of multifactor RNA-Seq experiments with respect to biological variation. Nucleic Acids Research 40, 4288–4297 (2012).

29. Robinson, M. D., McCarthy, D. J. & Smyth, G. K. edgeR: a Bioconductor package for differential expression analysis of digital gene expression data. Bioinformatics 26, 139–140 (2010).

30. Love, M. I., Huber, W. & Anders, S. Moderated estimation of fold change and dispersion for RNA-seq data with DESeq2. Genome Biology 15, 550 (2014).

31. van der Veeken, J. et al. Natural Genetic Variation Reveals Key Features of Epigenetic and Transcriptional Memory in Virus-Specific CD8 T Cells. Immunity 50, 1202-1217.e7 (2019).

32. Pritykin, Y. et al. A unified atlas of CD8 T cell dysfunctional states in cancer and infection. Molecular Cell 81, 2477-2493.e10 (2021).

33. Lin, Y. et al. Evaluating stably expressed genes in single cells. GigaScience 8, giz106 (2019).

34. Medina-Cano, D. et al. A Mouse Organoid Platform for Modeling Cerebral Cortex Development and Cis-Regulatory Evolution in Vitro. Preprint 2024.09.30.615887 (2024) doi:10.1101/2024.09.30.615887.

35. Metzger, B. P. H., Yuan, D. C., Gruber, J. D., Duveau, F. & Wittkopp, P. J. Selection on noise constrains variation in a eukaryotic promoter. Nature 521, 344–347 (2015).

36. Hounkpe, B. W., Chenou, F., de Lima, F. & De Paula, E. V. HRT Atlas v1.0 database: redefining human and mouse housekeeping genes and candidate reference transcripts by mining massive RNA-seq datasets. Nucleic Acids Res 49, D947–D955 (2021).

37. Eckersley-Maslin, M. A. et al. Random Monoallelic Gene Expression Increases upon Embryonic Stem Cell Differentiation. Developmental Cell 28, 351–365 (2014).

38. Arpi, M. N. T. & Simpson, T. I. SFARI genes and where to find them; modelling Autism Spectrum Disorder specific gene expression dysregulation with RNA-seq data. Sci Rep 12, 10158 (2022).

39. Collins, S. C. et al. Large-scale neuroanatomical study uncovers 198 gene associations in mouse brain morphogenesis. Nat Commun 10, (2019).

40. Naik, H. C. et al. Semicoordinated allelic-bursting shape dynamic random monoallelic expression in pregastrulation embryos. iScience 24, 102954 (2021).

41. Balasooriya, G. I. & Spector, D. L. Allele-specific differential regulation of monoallelically expressed autosomal genes in the cardiac lineage. Nat Commun 13, 5984 (2022).

42. Deng, Q., Ramsköld, D., Reinius, B. & Sandberg, R. Single-Cell RNA-Seq Reveals Dynamic, Random Monoallelic Gene Expression in Mammalian Cells. Science 343, 193–196 (2014).

43. Marion-Poll, L. et al. Locus specific epigenetic modalities of random allelic expression imbalance | Nature Communications. Nat Commun 12, (2021).

44. Gendrel, A.-V. et al. Developmental Dynamics and Disease Potential of Random Monoallelic Gene Expression. Developmental Cell 28, 366–380 (2014).

45. Kravitz, S. N. et al. Random allelic expression in the adult human body. Cell Reports 42, 111945 (2023).

46. Zhao, H. et al. Genome-wide fitness gene identification reveals Roquin as a potent suppressor of CD8 T cell expansion and anti-tumor immunity. Cell Reports 37, 110083 (2021).

47. Keane, T. M. et al. Mouse genomic variation and its effect on phenotypes and gene regulation. Nature 477, 289–294 (2011).

48. Danecek, P. et al. Twelve years of SAMtools and BCFtools. Gigascience 10, giab008 (2021).

49. Hinrichs, A. S. et al. The UCSC Genome Browser Database: update 2006. Nucleic Acids Res 34, D590–598 (2006).

50. Dobin, A. et al. STAR: ultrafast universal RNA-seq aligner. Bioinformatics 29, 15–21 (2013).

51. Griffiths, D. A. Maximum Likelihood Estimation for the Beta-Binomial Distribution and an Application to the Household Distribution of the Total Number of Cases of a Disease. Biometrics 29, 637–648 (1973).

52. R Core Team. R: A Language and Environment for Statistical Computing. (R Foundation for Statistical Computing, 2023).

53. Loader, C. Locfit: Local Regression, Likelihood and Density Estimation. (1999).

54. Hampel, F. R. The Influence Curve and Its Role in Robust Estimation. Journal of the American Statistical Association 69, 383–393 (1974).

55. Benjamini, Y. & Hochberg, Y. Controlling the False Discovery Rate: A Practical and Powerful Approach to Multiple Testing. Journal of the Royal Statistical Society 289–300 (1995).

56. McCarthy, D. J., Campbell, K. R., Lun, A. T. L. & Wills, Q. F. Scater: pre-processing, quality control, normalization and visualization of single-cell RNA-seq data in R. Bioinformatics 33, 1179– 1186 (2017).

57. Risso, D., Perraudeau, F., Gribkova, S., Dudoit, S. & Vert, J.-P. A general and flexible method for signal extraction from single-cell RNA-seq data. Nat Commun 9, (2018).

58. Yee, T. Vector Generalized Linear and Additive Models: With an Implementation in R | SpringerLink. (2015).

59. Hao, Y. et al. Integrated analysis of multimodal single-cell data. Cell 184, 3573-3587.e29 (2021).

60. Bolger, A. M., Lohse, M. & Usadel, B. Trimmomatic: a flexible trimmer for Illumina sequence data. Bioinformatics 30, 2114–2120 (2014).

61. Lilue, J., Shivalikanjli, A., Adams, D. J. & Keane, T. M. Mouse protein coding diversity: What’s left to discover? PLOS Genetics (2019).

62. Liao, Y., Smyth, G. K. & Shi, W. The R package Rsubread is easier, faster, cheaper and better for alignment and quantification of RNA sequencing reads. Nucleic Acids Res 47, e47 (2019).

63. Kaminow, B., Yunusov, D. & Dobin, A. STARsolo: accurate, fast and versatile mapping/quantification of single-cell and single-nucleus RNA-seq data. 2021.05.05.442755 Preprint at (2021).

64. Lun, A. T. L., McCarthy, D. J. & Marioni, J. C. A step-by-step workflow for low-level analysis of single-cell RNA-seq data with Bioconductor. F1000Res 5, 2122 (2016).

65. Grant, C. E., Bailey, T. L. & Noble, W. S. FIMO: scanning for occurrences of a given motif. Bioinformatics 27, 1017–1018 (2011).

66. Quinlan, A. R. & Hall, I. M. BEDTools: a flexible suite of utilities for comparing genomic features. Bioinformatics 26, 841–842 (2010).

67. Ho, D., Imai, K., King, G. & Stuart, E. A. MatchIt: Nonparametric Preprocessing for Parametric Causal Inference. Journal of Statistical Software 42, 1–28 (2011).

68. Wu, T. et al. clusterProfiler 4.0: A universal enrichment tool for interpreting omics data. Innovation (Camb) 2, 100141 (2021).

69. Carlson, M. & Falcon, S. Org. Mm. Eg. Db: Genome Wide Annotation for Mouse. vol. 3 (R package, 2019).

70. Yang, M. G., Ling, E., Cowley, C. J., Greenberg, M. E. & Vierbuchen, T. Characterization of sequence determinants of enhancer function using natural genetic variation. eLife 11, e76500 (2022).

