## Supplemental Figures for "ASPEN: Robust detection of allelic dynamics in single cell RNA-seq"

[Supplemental Figure 8. Selecting appropriate number of permutations with tolerance < 0.01.	13](#_Toc205467213)

### Supplemental Results

**Result of including multiple mapping reads**

We tested the inclusion of weighted multimapping reads using STARSolo’s EM method to probabilistically assign multimappers based on gene expressed across all cells (Kaminow, 2021)(**Methods**). Inclusion of multimapping reads resulted in recovery of ~2-fold of allelic reads, from 26.7M to 47.2M on average, based on the B6xSpret F1 data (**Supp Fig. 10A**). However, most of those reads do not overlap an informative variant. We detected increased dispersion (**Supp Fig. 10B**). Despite evaluating a greater number of genes when using both the multimapping and unique genes, 27.7% fewer genes had significant allelic imbalance (ASPEN-mean FDR < 0.05, n=1,167) than those identified using unique counts (**Supp Fig. 10C**). Our observations suggested that the inclusion of multi-mapping reads leads to lower sensitivity in allelic imbalance detection.

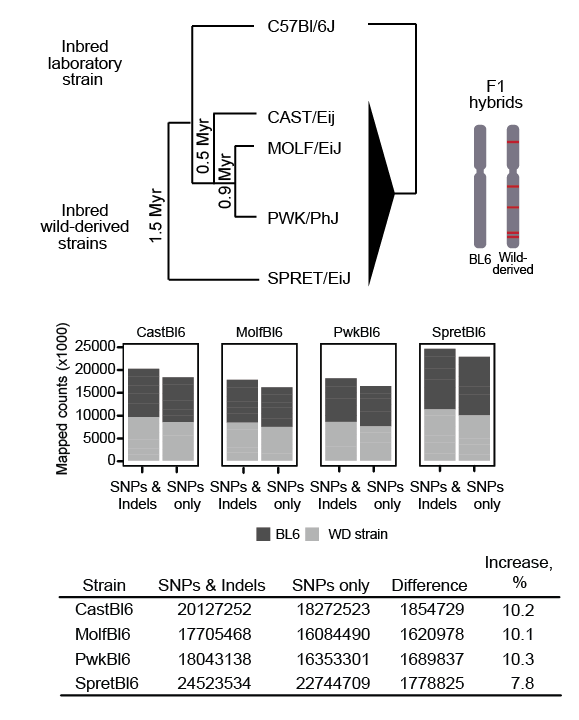

### Supplemental Figure 1. **Comparing two F1 hybrid reads mapping strategies: SNPs level variation and SNPs and indels level variation between the parental genomes.**

Schematic for F1 breeding strategy (top panel). The difference in the number of reads mapped to the combined genomes generated with either SNPs only or both SNPs and indels using fibroblast cells from four mouse F1 hybrids – Bl6 × Cast, Bl6 × Molf, Bl6 × Pwk and Bl6 × Spret (bottom panel).

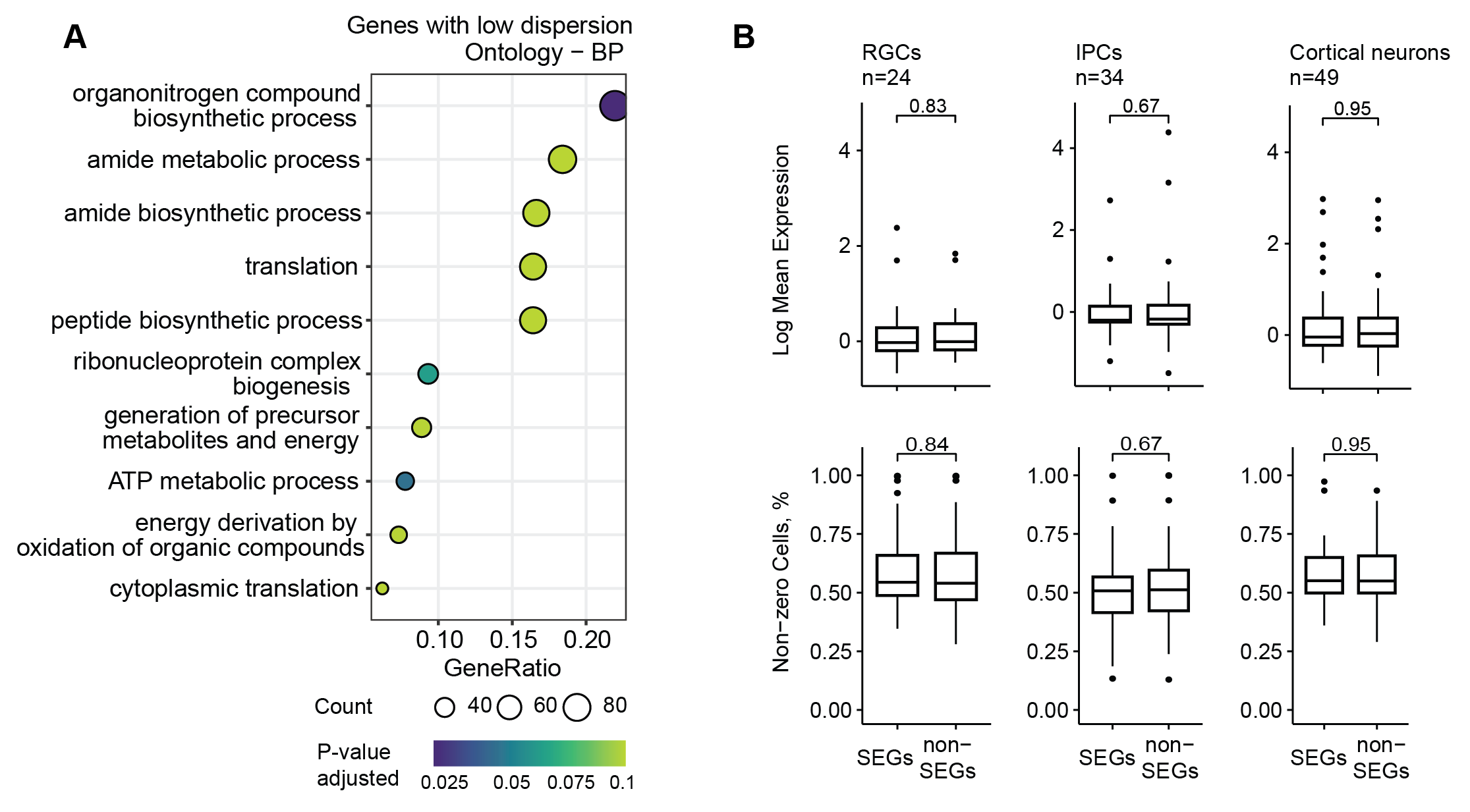

### **Supplemental Figure 2. Genes with low dispersion involved in broad cellular functions.**

**(A)** GO term enrichment analysis using the genes with low dispersion. Enriched gene ontology biological process function categories for genes with low allelic dispersion in T-cells day 7 post LCMV infection scRNA dataset **(B)** Mean expression and proportion of non-zero cells between the stably expressed genes (SEGs), identified in Bl6 × Cast F1 brain organoids dataset, and background, genes matched by expression level and number of non-zero cells (two-sided Wilcoxon rank-sum test) (complementary to Fig. 2C).

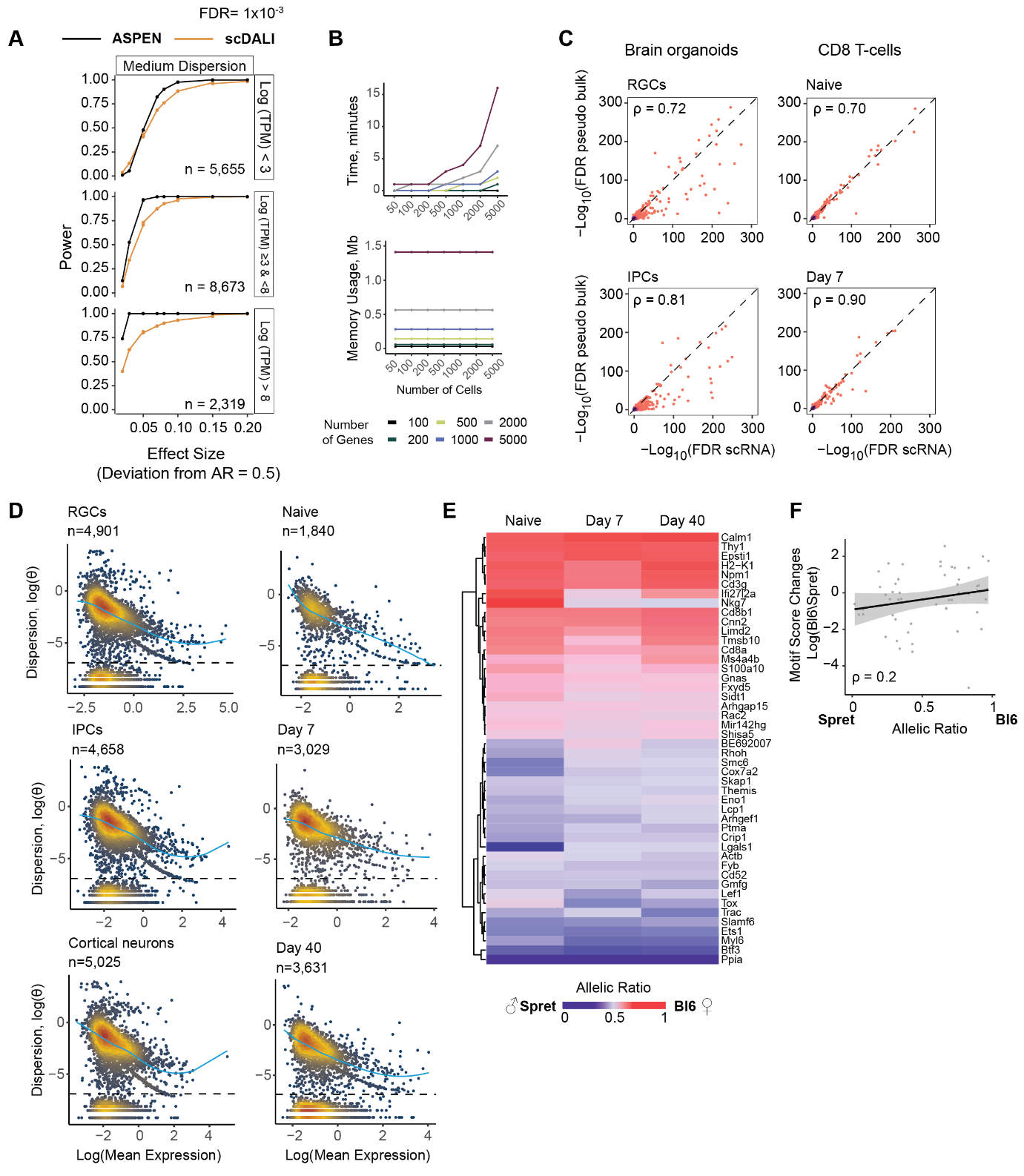

### **Supplemental Figure 3. Detecting genes with allelic imbalance using ASPEN.**

**(A)** Performance comparison between ASPEN and scDALI using simulated data (complementary to Figure 3C). Line plots showing TPRs calculated at FDR = 0.001 for genes stratified by gene expression and allelic dispersion levels (low dispersion – θ < 0.005, medium – θ ≥ 0.005 & θ < 0.4 and high – θ ≥ 0.4) across groups with varying levels of deviation from the mean AR = 0.5. Only datasets simulated with a mean AR of 0.5 $\pm$ 0.2 are shown. **(B)** ASPEN runtime (top) and memory usage (bottom) for different numbers of cells and genes (calculations were restricted to one core using a 6 CPU Intel Xeon, 64GB RAM machine). **(C)** Correlation of -log_10_FDR values from ASPEN mean test ran on the single-cell counts (x-axis) and pseudobulked counts (y-axis) (complementary to Figure 3E). Analyses were performed in mouse brain organoid cells from female Bl6 × Cast F1 hybrids (left panel): radial glial cells (RGCs, n = 1,072) and intermediate progenitors (IPCs, n = 853); and in CD8^+^ T cells from male Bl6 × Spret F1 hybrids (right panel): the naïve state (n = 221) and day 7 (n = 525) after LCMV infection. **(D)** Relationship between allelic ratios dispersion and mean gene expression in organoid cells from female Bl6 × Cast F1 hybrid (left panel) and CD8 T-cells from male Bl6 × Spret F1 hybrid (right panel). The dashed line separates genes with low variation which are excluded from the shrinkage procedure. The local regression model fit is shown in light blue. **(E)** Top 50 genes with significant allelic imbalance (ASPEN mean FDR < 0.05) identified in T-cells from male Bl6 × Spret F1 hybrids and consistently expressed in all three cell states. **(F)** Correlation between allelic ratio and changes in motif score for the variants inside the promoter peaks (+/- 1Kb from TSS). Allelically imbalanced genes with strong bias to either Bl6 or Spret allele are selected (ASPEN-mean FDR < 0.05, |log2FC| $\geq$ 1).

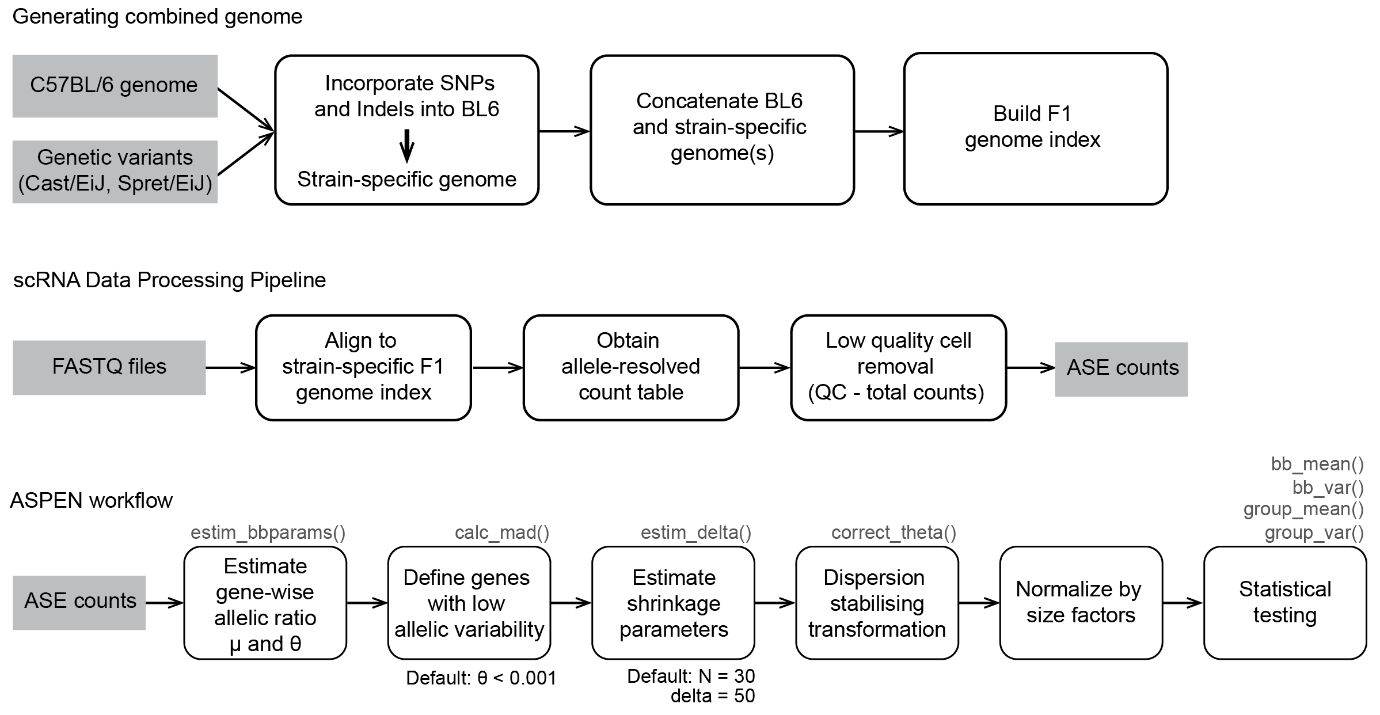

### **Supplemental Figure 4. ASPEN workflow**.

scRNA data pre-processing steps, including the generation of the F1 index file for ASE reads mapping (top row) and ASPEN pipeline (bottom row). Tests evaluating allelic variation (deviation from the expected level of dispersion for genes with similar expression – bb_var, allelic variance changes across the groups – group_var) are performed using raw counts. For the group-level tests (group_mean and group_var), dispersion stabilization steps are to be repeated for each group and across all cells. The counts normalization is performed outside of ASPEN. We used computeSumFactors from scran (Lun., et al. 2016). The minimum coverage threshold for the genes is set to a minimum of five reads in at least five cells (min_counts = 5, min_cells = 5). We find this to be sufficient to reliably identify ASE patterns in most datasets, however for the differential changes in variance we recommend increasing the minimum number of cells to 15 (min_cells = 15).

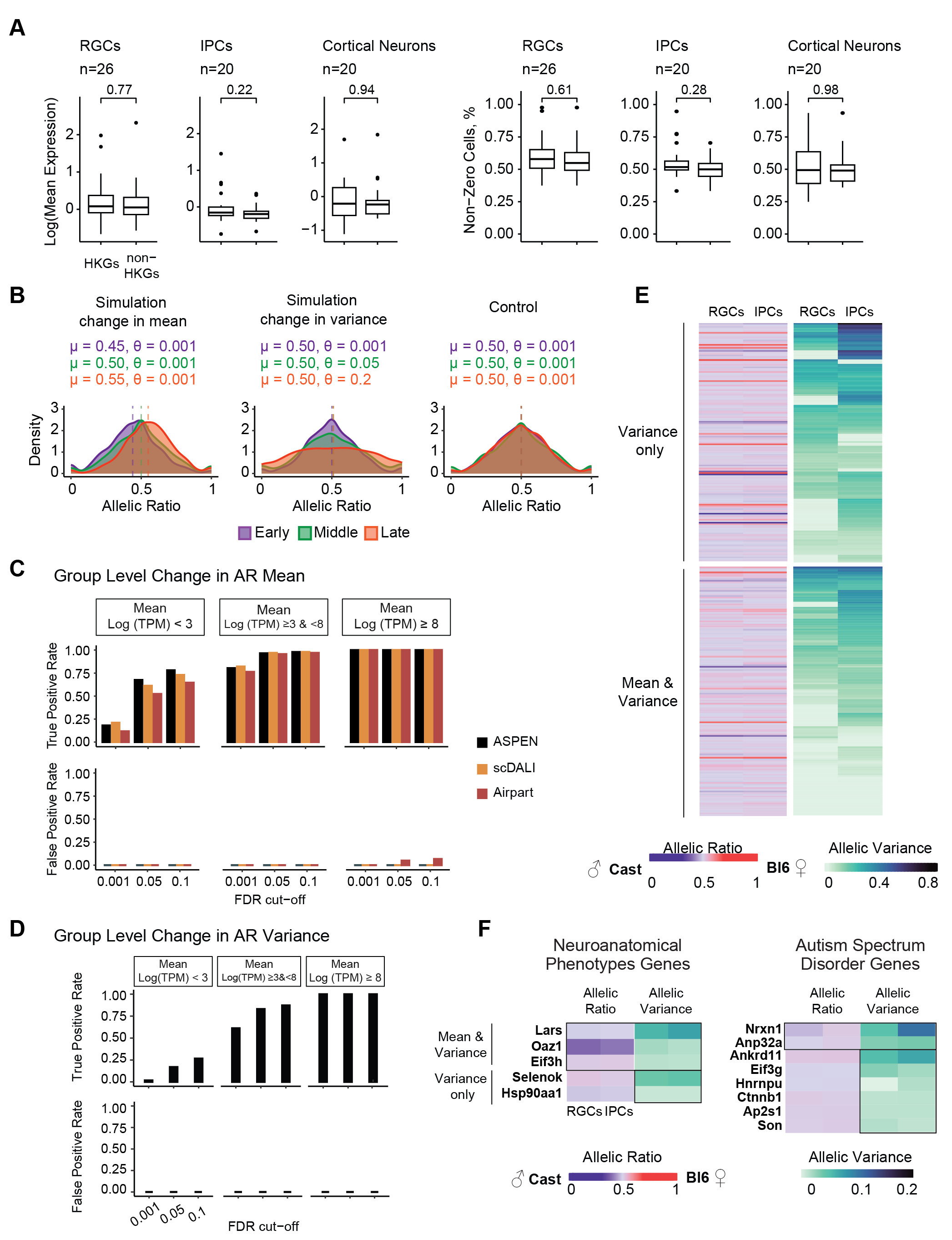

### **Supplemental Figure 5. Detecting group-level changes in allelic patterns during early neurogenesis.**

**(A)** Mean expression and proportion of non-zero cells between the housekeeping genes (HKGs), identified in Bl6 × Cast F1 brain organoids dataset, and background, genes matched by expression level and number of non-zero cells (two-sided Wilcoxon rank-sum test) (complementary to Fig. 4D). **(B)** Allelic ratio distribution in three simulated scenarios: changes in mean AR (with dispersion parameters held constant), changes in AR variance (with mean parameters held constant), and a control group (where both mean and dispersion parameters are constant). The reference allele counts were simulated by drawing from a beta-binomial distribution parameterized by the bin-wise total counts, mean (μ), and dispersion (θ) for the respective bin as indicated. **(C)** Comparing the accuracy in detecting allelic ratio change across the groups between ASPEN, scDALI and Airpart in different gene expression groups. **(D)** Accuracy in detecting changes in allelic variation improves as the number of informative cells with sufficient coverage increases. **(E)** Global overview of the genes with group-level allelic variation in differentiation from radial glial cells (RGCs) to intermediate progenitor cells (IPCs) in female Bl6 x Cast F1 hybrids. Genes are partitioned by changes in variance (n = 158) and both mean and variance (n = 148). **(F)** Genes with differential allelic patterns between RGCs and IPCs are associated with either Intellectual Disability/Autism Spectrum Disorders (ID/ASD, SFARI score 1) or with neuroanatomical phenotypes in mice.

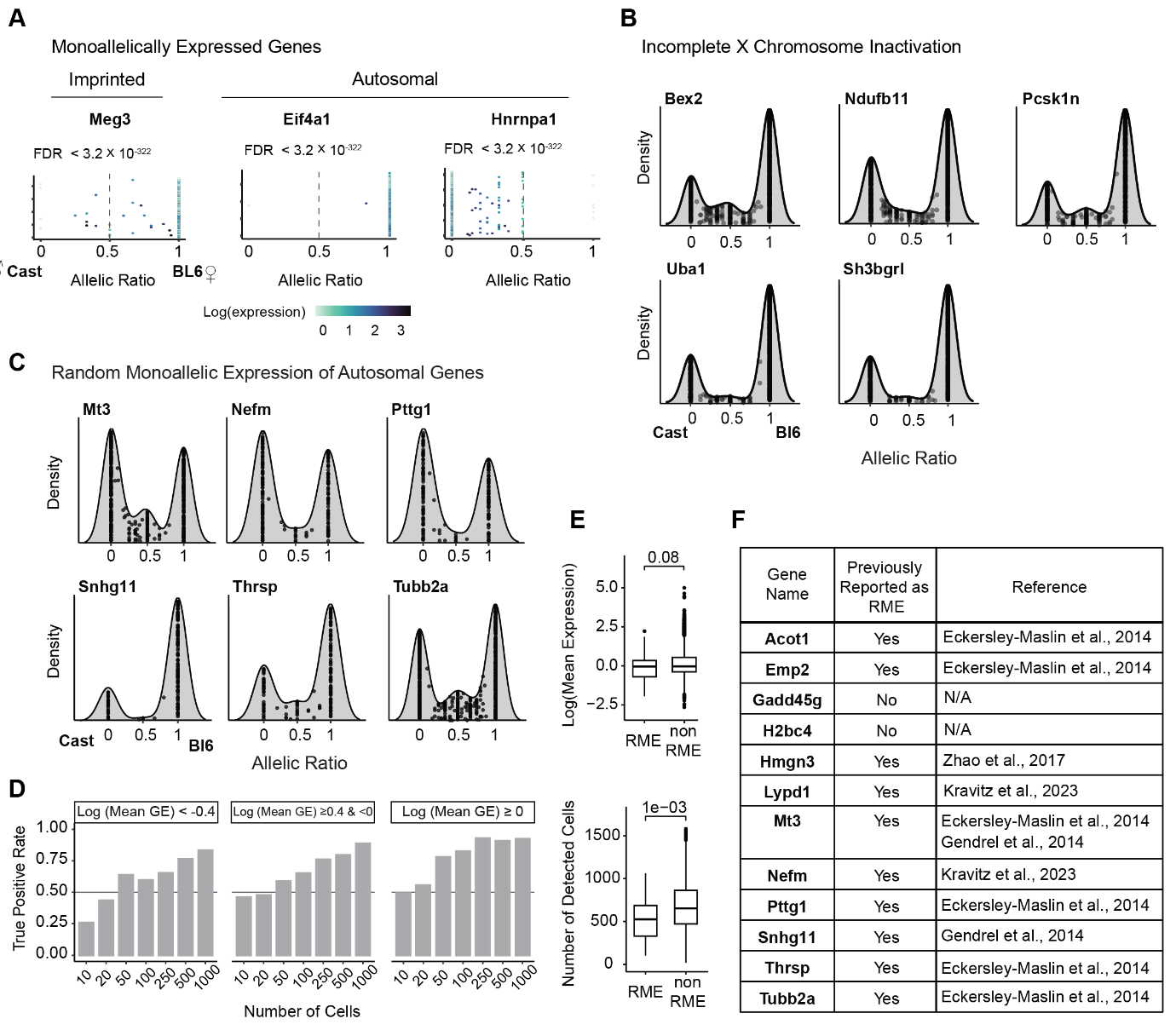

### **Supplemental Figure 6. Evaluating monoallelic ASE patterns in mouse brain organoids from female Bl6 x Cast hybrid**.

**(A)** Examples of AR distribution in genes with monoallelic expression (defined using beta-binomial shape parameters α < 1 or β < 1). Genes showing consistent monoallelic expression on either of the alleles include imprinted and autosomal genes. FDR values indicate deviation from balanced ASE (ASPEN mean test). **(B)** Allelic ratio distribution of genes with incomplete X chromosome inactivation in female Bl6 × Cast F1 hybrid. **(C)** Allelic ratio distribution plots for the autosomal genes with random monoallelic expression (RME) found in the brain organoid data (complementary to Figure 5D). **(D)** True positive rate for RME detection in the simulated data. 500 genes were simulated with different numbers of cells (n = 10, 20, 50, 100, 250, 500, 1000) and stratified by gene expression. **(E)** Median expression and median number of non-zero cells between RME and non-RME genes. Median expression: RME – 0.96, non-RME – 0.97, D = 0.182, p = 0.08, two-sided K-S test. Median number of cells: RME - 526, non RME – 654, D = 0.279, p = 0.001, two-sided K-S test. **(F)** List of autosomal genes identified as RME (defined using beta-binomial shape parameters α < 1 and β < 1 and |α – β| < 0.5) with evidence of previous reports in the literature.

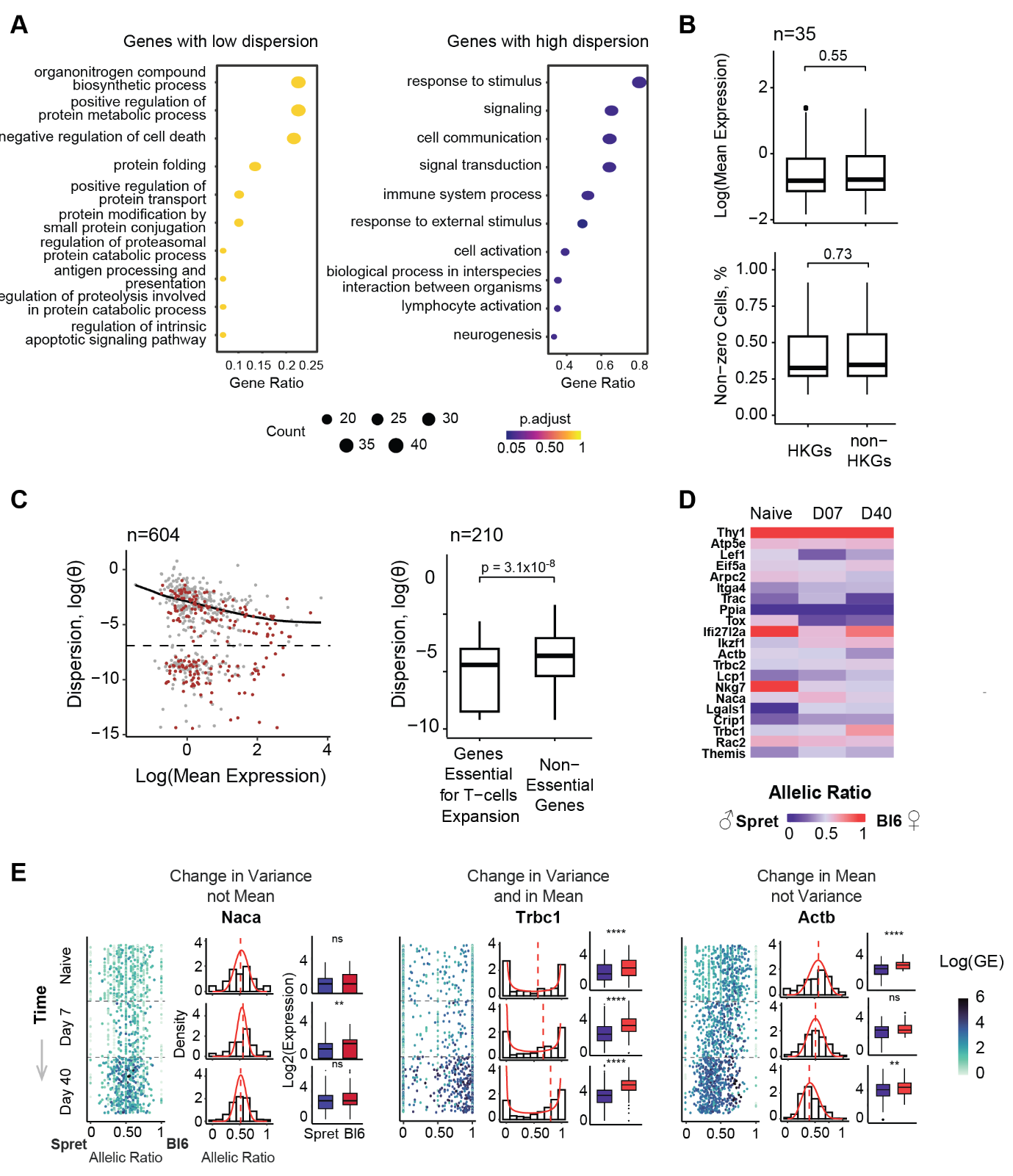

### Supplemental Figure 7. Allelic variation patterns during T-cell activation.

**(A)** Gene Ontology (GO) term enrichment analysis result for genes with allelic variance different from the expected for similar expression levels (ASPEN var: FDR < 0.05). The analysis included genes across all three states (naïve, day 7 and day 40). The genes were divided into low-dispersed (empirical dispersion is less than common) and highly dispersed (empirical dispersion is greater than common) groups. **(B)** Mean expression and proportion of non-zero cells between the housekeeping genes, identified in low-dispersed T cells, and background, genes matched by expression level and number of non-zero cells (two-sided Wilcoxon rank-sum test) (complementary to Fig. 6B). **(C)** Abundance of genes essential for the T-cells expansion (marked in red) in day 7 post LCMV infection scRNA dataset. The boxplot shows mean dispersion estimation between essential genes and background, a set of genes matched by gene expression (two-sided Wilcoxon rank-sum test). **(D)** Mean allelic ratio across the cell states for genes with significant deviation from allelic trend (complementary for Fig. 6B) **(E)** Boxplots showing the mean ASE (two-sided Wilcoxon rank-sum test, ‘****’ *p* ≤ 1 × 10^-4^, ‘**’ *p* ≤ 1 × 10^-2^) for genes with varying temporal distributions: *Naca* (changes in allelic variance), *Actb* (changes in allelic mean) and *Trbc1* (changes in both allelic variance and mean) (complementary to Fig. 6E).

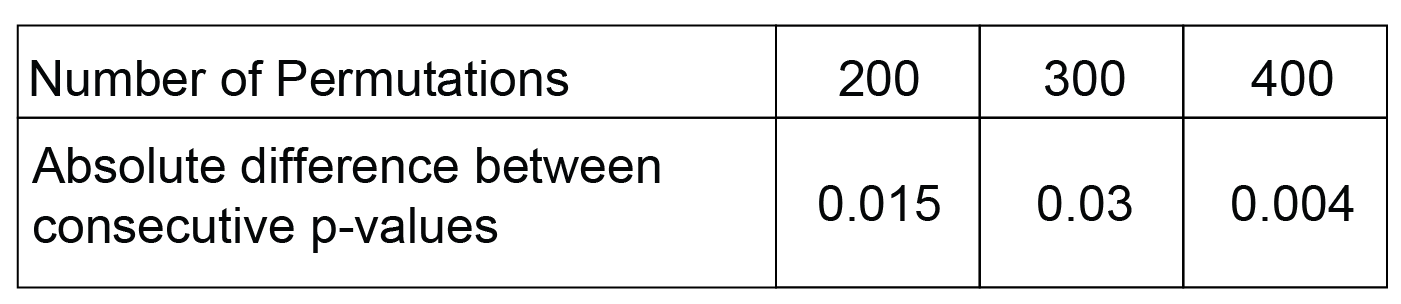

### Supplemental Figure 8. Selecting appropriate number of permutations with tolerance < 0.01.

We used data simulated with mean AR = 0.42 and tested different number of permutations (100 for each step) to calculate the *p*-value difference for one point. A tolerance below 0.01 was reached after 400 permutations.

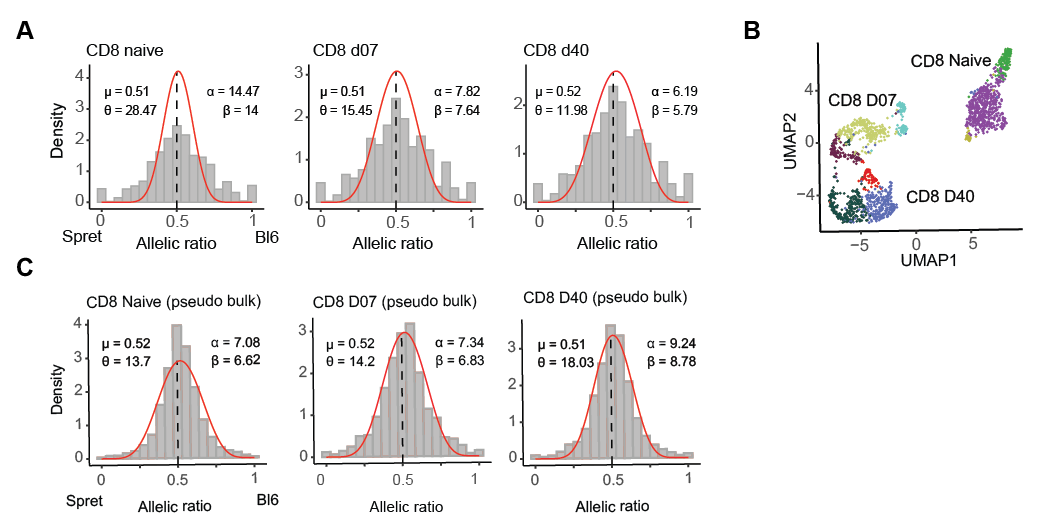

### **Supplemental Figure 9. Allelic imbalance analyses using single-cell and pseudobulked counts in mouse T-cells from male Bl6 x Spret hybrid.**

**(A)** Global AR distribution across all genes in the T cells dataset (scRNA counts), showing a slight bias at 0.52 towards the BL6 allele. **(B)** UMAP showing clustering of the T cells within each cell state that was used for the pseudo-bulk aggregation **(C)** Global AR distribution across all genes in the mouse T-cells dataset (male Bl6 x Spret hybrid, using pseudo-bulked scRNA counts), showing a slight bias towards the BL6 allele.

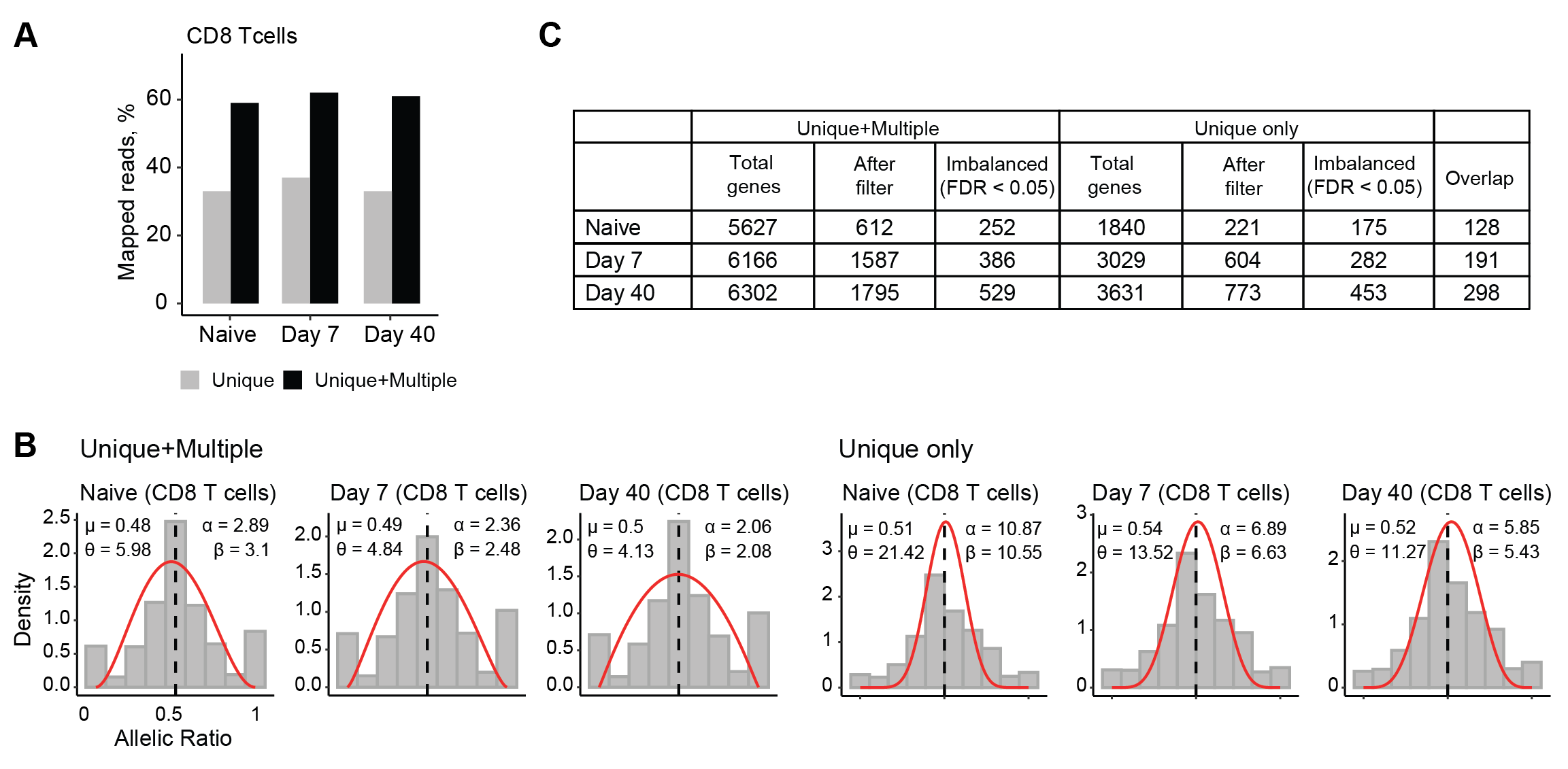

### Supplemental Figure 10. Expectation Maximization (EM) strategy for the multimapping reads inflates the number of genes with low expression.

**(A)** Inclusion of multi-mappers leads to ~2-fold increase in the number of allelically resolved reads. Using the T-cell data, the reads were mapped against the Bl6 × Spret F1 genome. The average increase in the number of mapped reads was from 34.3% (unique) to 60.6% (unique + multiple). **(B)** Global AR distribution in genes quantified using multimapping and unique reads (top row) and unique reads only (bottom row). Using multi-mappers for gene quantification leads to higher dispersion around the mean AR. **(C)** ASPEN-mean test results for two gene categories: including the multimapping reads and using the unique reads only. The total number of genes (after quantification), the number of genes after applying the minimum coverage threshold (at least five cells with a minimum of five reads), the number of genes with allelic imbalance (ASPEN-mean FDR < 0.05) and the overlap between the two groups are shown. A smaller fraction of allelically imbalanced genes in the first gene category suggests that using multimapping reads does not necessarily lead to better accuracy in detecting allelic imbalance.
